# Surrogate production of genome edited sperm from a different subfamily by spermatogonial stem cell transplantation

**DOI:** 10.1101/2021.04.20.440715

**Authors:** Fenghua Zhang, Xianmei Li, Yongkang Hao, Yi Li, Ding Ye, Mudan He, Houpeng Wang, Zuoyan Zhu, Yonghua Sun

## Abstract

The surrogate reproduction technique provides a powerful tool for production of allogenic or xenogeneic gametes derived from endangered species or those with valuable genetic traits. Production of functional donor-derived gametes through intra- or inter-specific spermatogonial stem cell transplantation (SSCT) has been achieved in many species. However, generation of functional gametes from a phylogenetically distant species such as from a different subfamily by SSCT has never been successful. Here, using two small cyprinid fishes, Chinese rare minnow (*gobiocypris rarus*, for brief: *Gr*) and zebrafish (*danio rerio*), which belong to different subfamilies, as donors and recipients for SSCT, we optimized the SSCT technique and successfully obtained *Gr*-derived sperm carrying targeted genome modifications in zebrafish. We revealed that the transplanted *Gr* spermatogonia supported the host gonadal development and underwent normal spermatogenesis, resulting in a reconstructed fertile testis containing *Gr* spermatids and zebrafish testicular somatic cells. Interestingly, the surrogate spermatozoa resembled those of host zebrafish but not donor *Gr* in morphology and swimming behavior. Finally, we showed that *Gr*-derived genome edited sperm was successfully produced in zebrafish by cross-subfamily SSCT, when the *pou5f3* and *chd* gene knockout *Gr* SSCs were used as surrogate donors. This is the first report demonstrating the surrogate production of genome edited sperm from a phylogenetically distant species, and this method is feasible to be applied to future breeding of commercial fishes.

## 1. Introduction

As a unique cell type that can transmit genetic materials to next generation, germ cells are capable of generating an entire organism and maintaining the everlasting germline cycle therefore immortal (Cinalli et al., 2008). Germline stem cells (GSCs) are considered as a type of undifferentiated germ cells with the ability of self-renewal, and GSC transplantation (GSCT) technology provides a powerful tool to obtain the gametes with the genetic properties of donor cells from surrogates (Lehmann, 2012). The GSCT was first reported in domestic chicken by transferring the isolated primordial germ cells (PGCs) into the blood stream of embryos (Tajima et al., 1993), and later in mice by transplanting the spermatogonial stem cells (SSCs) (Brinster, 2002). In fish, this technology was first established in zebrafish (Ciruna et al., 2002), a widely used model animal, and then expanded to some economically commercial fishes like salmonids (Takeuchi et al., 2004) and rainbow trout (Okutsu et al., 2006). In previous experiments, PGCs, SSCs and ovogenic stem cells (OSCs) have been used as donors for transplantation into embryos, larvae or adults of another species (Goto and Saito, 2019). Among all types of donor GSCs, SSCs are most extensively studied and thus SSCT is most widely utilized for surrogate reproduction (Kubota and Brinster, 2018). It is proposed that the application of xenogeneic SSCT is promising in broodstock breeding, protection of rare and endangered species and sex control breeding.

The clustered regularly interspaced short palindromic repeats (CRISPR)/CRISPR-associated 9 (Cas9) technology has been used in several fish species to study gene functions or to potentially improve aquaculture breeding, and a large amount of genetic-null mutants have been generated in model species such as zebrafish (*Danio rerio*, subfamily *danioninae*) (Chang et al., 2013; Sun et al., 2020). However, when the CRISPR/Cas9 technology is used for generating genetically homozygous mutants of a commercial fish with relative long breeding cycle, it will be a time-and labor-consuming process. In addition, if we intend to generate genome edited fish with large body size, large and expensive facilities are normally necessary for rearing and screening the individuals. Recently we developed a novel approach to generate zebrafish maternal-zygotic mutants of embryonic lethal genes in a relative short period, by combining the mediated genome editing technology and the optimized PGCs transplantation (PGCT) technique (Zhang et al., 2020). However, it remains unknown whether genome edited germ cells could be generated by surrogate reproduction between two phylogenetically distant species, in order to utilize this technology to expedite genetic breeding in aquaculture.

Chinese rare minnow, (*Gobiocypris rarus*, subfamily *gobioninae*, abbr: *Gr*), is a small cyprinid fish that distributes specifically in tributaries of the Dadu River in the upper reaches of the Yangtze River and some small rivers in Sichuan province, China. Recently, *Gr* can rarely be seen in wild environment because of habitat destruction, introduction of carnivorous exotic fishes and overfishing, and it has already been listed as endangered in China Species Red List (Luo et al., 2017). In recent years, this species has been developed as a laboratory fish to study ichthyopathology, genetics, environmental science, embryology and physiological ecology, owning to its qualified characteristics, such as small size, ease of culture, high fecundity, transparent embryos and sensitivity to toxicants (Liang and Zha, 2016). Different from zebrafish that reaches sex maturity as early as 2.5 months post fertilization (mpf) (Zhang et al., 2020) and spawns in the morning (Eaton and Farley, 1974), *Gr* only matures at 4 mpf and spawns during 18:00 to 24:00 pm (Wang, 1992). Thus, these two laboratory fish species from different subfamilies are ideal for exploring the feasibility of surrogate production of evolutionarily distant species derived genome edited gametes, by combination of CRISPR/Cas9 technology and xenogeneic SSCT.

Here, using zebrafish as surrogates and *Gr* as germline donor, we optimized the SSCT procedure and successfully obtained functional genome edited *Gr* sperm in zebrafish in less than 3 months. We clearly clarified the dynamic process, namely the colonization, proliferation, and differentiation, of *Gr* germ cells in the zebrafish surrogates. Intriguingly, the *Gr*-derived donor sperm produced by zebrafish hosts resembled the host sperm rather than the donor sperm in morphology and swimming behavior.

## 2. Results

### 2.1 Purification of *Gr* SSCs and SSCT into zebrafish larvae

The procedure for CRISPR/Cas9 induced xenogeneic SSCT were shown in Figure 1A. Briefly, *Gr* testes were dissected and enzymatically digested for purification of SSCs by Percoll gradient as described (Yoshikawa et al., 2009). The purified SSCs were transplanted into the hatching zebrafish larvae deprived of endogenous PGCs by *dead end* (*dnd*) antisense morpholino (MO) injection as previously described (Weidinger et al., 2003; Zhang et al., 2020). To determine which layer contained the richest SSCs after Percoll purification, we checked the interfaces from 25% to 60% and found that SSCs emerged in fractions of 25% to 40% with the richest in 35%, while 50% and 60% consisted almost exclusively spermatids and spermatozoa (Figure 1B). Number of SSCs was calculated by a cell counting chamber and about 1.45×10^4^ and 6.4×10^3^ SSCs per fish could be harvested in fractions of 35% and 40% respectively (Figure 1C). The purified SSCs were confirmed by its nuclear diameter of 4.7-8.6 μm (Leal et al., 2009) and by the expression of a SSCs-specific marker, Nanos2 (Beer and Draper, 2013; Bellaiche et al., 2014) (Figure 1D). Therefore, these two cell fractions were collected for SSCT. Similar to our previous study (Zhang et al., 2020), when using *Gr pou5f3* and *chd* knocked out (KO) adult males as donors, the mutation efficiencies of these two genes in isolated SSCs reached as high as 100.0% and 96.7% respectively (Figure S1). To visualize the donor SSCs in vivo, we stained them with live cell tracker of red fluorescence (Figure 1E, arrows) and transplanted them into the peritoneal cavity of sterile zebrafish larvae between the swim bladder and gut close to the primitive gonads (Figure 1F). Three days post transplantation (dpt), large numbers of labeled *Gr* donor cells were observed in the primitive gonad region of host zebrafish, indicating that *Gr* donor SSCs survived and colonized in zebrafish surrogates (Figure 1G, arrows).

**Figure 1.**
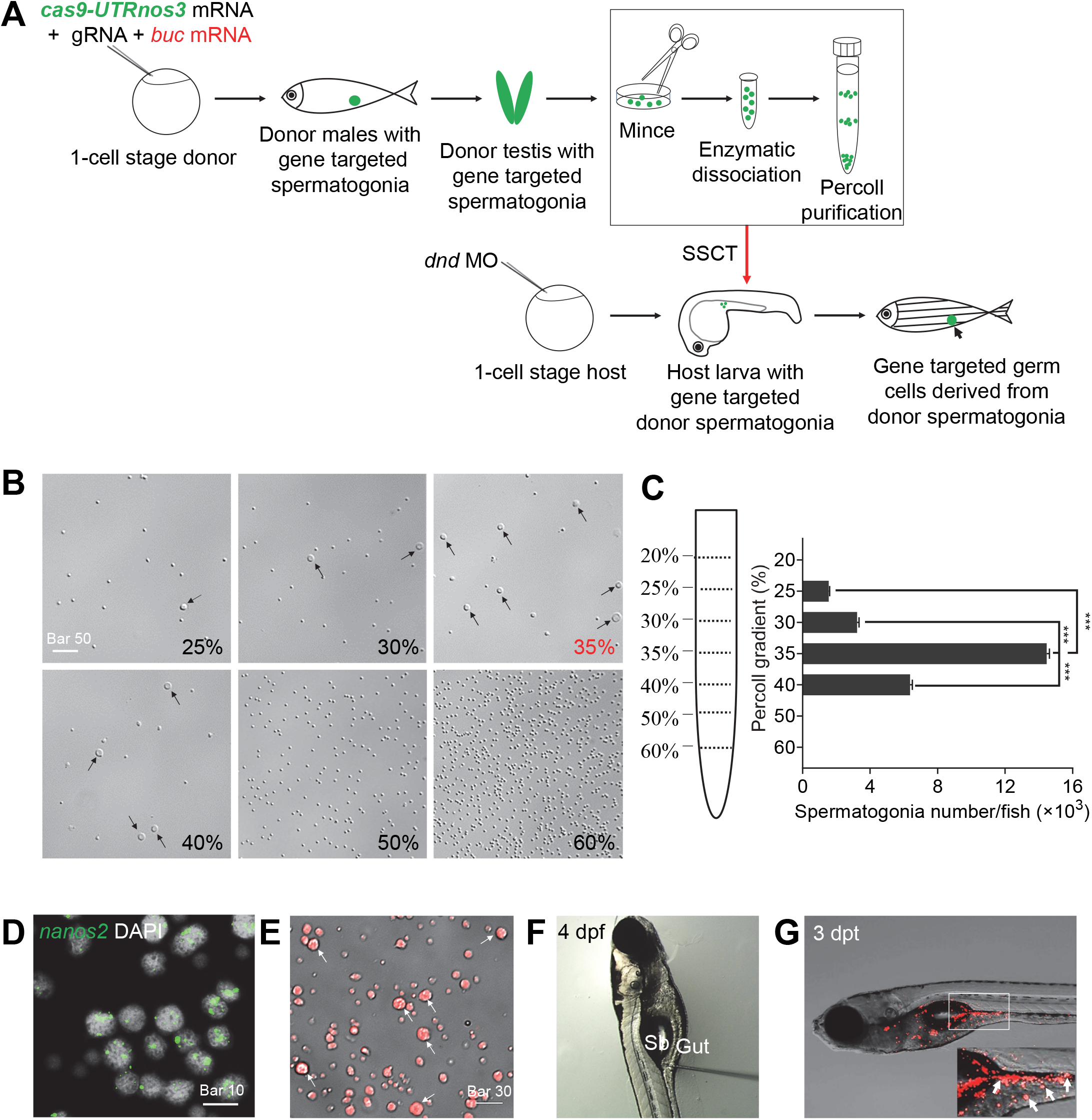
Spermatogonia stem cells purification and transplantation. **A**: Schematic workflow represents process of the optimized procedure of SSCs-targeted CRISPR/Cas9 and transplantation of purified SSCs between different fish species. **B**: Germ cells purified from different Percoll gradient fractions (25%, 30%, 35%, 40%, 50%, and 60%). Arrows showed the SSCs in different fractions. Note that the most SSCs could be observed at 35% fraction. **C**: The number of SSCs per fish obtained from each gradient fraction (mean ± SEM). **D**: Fluorescent in situ hybridization by *Gr nanos2* probe indicated that the purified cells with about 5-9 μm nuclear diameter were spermatogonia stem cells. Scale bar, 10 μm. **E**: Viable donor cells were labeled with red fluorescent dye and the arrows showed the donor SSCs. Scale bar, 30 μm. **F**: Purified and labeled SSCs were transplanted into the abdominal cavity between the swim bladder (Sb) and gut of 4 dpf sterile zebrafish larva. **G**: Fluorescent image of recipient larvae at 3 days post transplantation (dpt). Arrowheads indicate numerous red fluorescence labeled donor cells in the genital region of the recipient larvae. Abbreviations: SSCs, spermatogonia stem cells; SEM, standard error of mean; *Gr, gobiocypris rarus*.

### 2.2 Optimization of donors and hosts for SSCT between *Gr* and zebrafish

Our previous work demonstrated that *bucky ball* (*buc*) mRNA overexpression could induce ectopic PGCs in zebrafish embryos, and significantly improved success rate of allogenic PGCs transplantation (PGCT) (Zhang et al., 2020). Since PGCs are embryonic progenitors of SSCs (Ohta et al., 2004), we tested whether overexpression of zebrafish *buc* (*zbuc*) could also increase SSCs number in *Gr* and elevate the success rate of xenogeneic SSCT. We injected *zbuc* mRNA into 1-cell stage *Gr* embryos at a dosage of 200 pg per embryo, and found that the PGCs were largely increased at blastula stage compared to uninjected wild type (WT) embryos, as visualized by whole-mount in situ hybridization of *vasa* (Figure 2A, B). Furthermore, as visualized by *GFP-UTRnos3* mRNA injection (Saito et al., 2006), overexpression of *zbuc* could persistently induce additional PGCs in 2 dpf *Gr* larvae (Figure 2C, D). Just like previously reported (Slanchev et al., 2005; Weidinger et al., 2003), all the *dnd* MO injected zebrafish grew to be infertile males. However, some *dnd* morphant zebrafish surrogates recovered their fertility after receiving xenogeneic *Gr* SSCs (Figure 2E), indicating the success of SSCT. Finally, we showed that the success rate of SSCT was significantly increased when using *zbuc*-overexpressed *Gr* males as the donors (Figure 2E).

**Figure 2.**
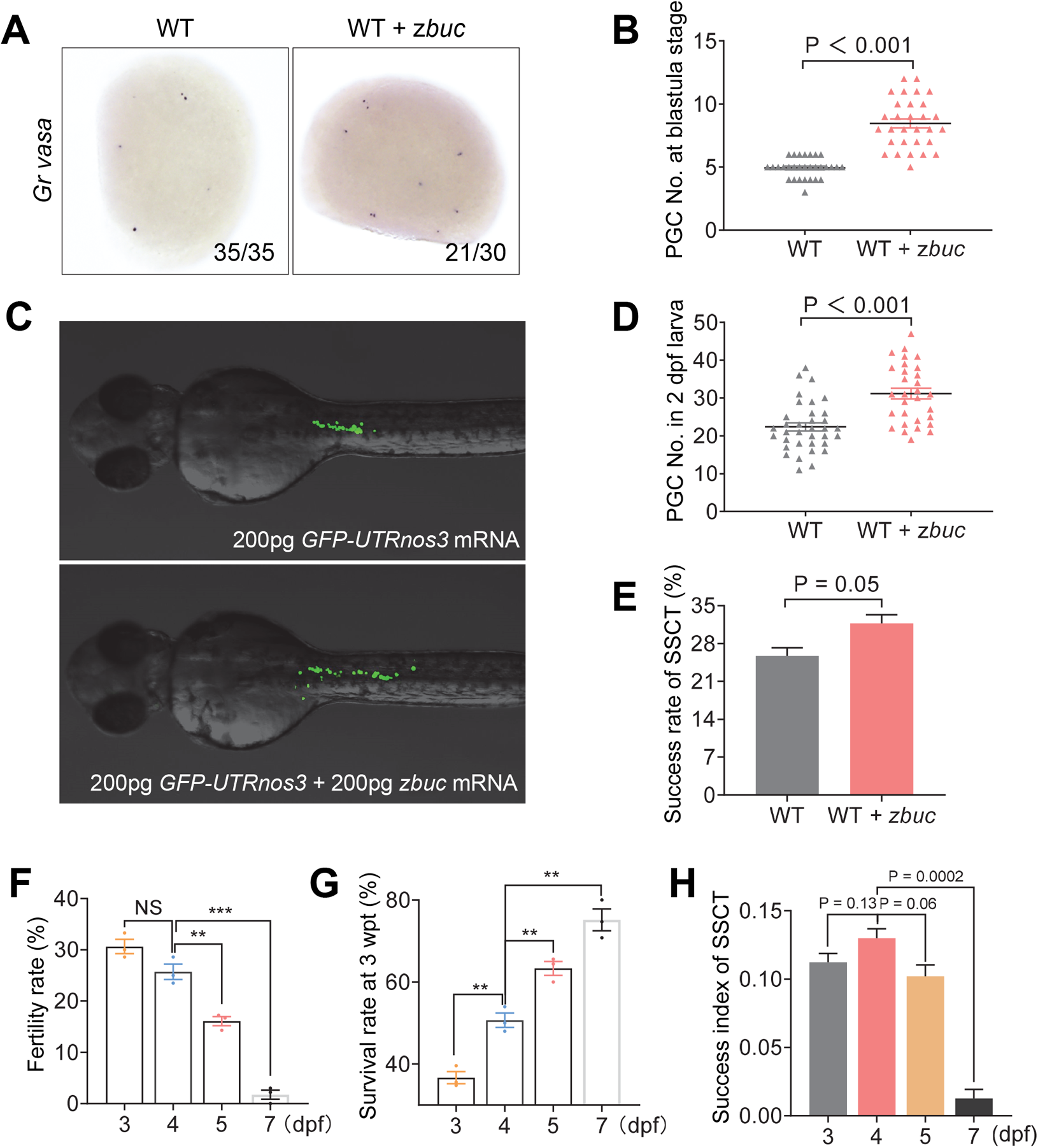
Optimization of donor and host for SSCT between *gobiocypris rarus* and zebrafish. **A**: Whole-mount in situ hybridization with *Gr vasa* probe of the wild type (WT) *Gr* blastula embryos (WT) and the blastula embryos injected with 200 pg *zbuc-UTRsv40* mRNA at 1-cell stage (WT + *zbuc*). The embryos were photographed from animal pole view. Note that 8-11 *vasa* positive cells appeared in the *zbuc*-injected embryos, compared with 5 *vasa* positive cells in the control group. **B**: Statistics of PGCs number of *Gr* embryos at blastula stage show that, compared with control group, the embryos injected with *zbuc* mRNA contain much more PGCs. **C**: 200 pg *zbuc* mRNA injection at 1-cell stage induced ectopic PGCs in *Gr* embryos at 2 dpf. **D**: Statistics of PGCs number in *Gr* embryos at 2 dpf show that, compared with control embryos, the embryos injected with *zbuc* mRNA contain much more PGCs. **E**: The success rate of SSCT, as indicated by fertility rate at adult stage, was increased by injection of *zbuc* mRNA into *Gr* donor embryos, utilizing 4 dpf zebrafish as host. The experiment was replicated for three times. **F**: The fertility rate of SSCT adults when 3, 4, 5, 7 dpf zebrafish larvae were utilized as hosts. **G**: The survival rate of SSCT individuals at 3 wpt when 3, 4, 5, 7 dpf zebrafish larvae were utilized as hosts. **H**: The success index of SSCT (survival rate at 3 wpt × fertility rate at adult stage) with different stages of host larvae at 3,4,5,7 dpf. It was the highest when 4 dpf zebrafish larvae were used as recipients. The experiment was replicated for three times. Abbreviations: SSCT, spermatogonial stem cell transplantation; *Gr, gobiocypris rarus*; PGCs, primordial germ cells; dpf, days post fertilization; wpt, weeks post transplantation. *zbuc*, zebrafish *bucky ball*.

To optimize the stage of host zebrafish embryo for transplantation of *Gr* SSCs, 3, 4, 5, 7 dpf zebrafish larvae were used (Figure S2), and the fertility rate (number of fish could fertilize WT eggs / number of fish tested) of the SSCT adults were compared. The fertility rate of survival SSCT adults with host larvae at 3 dpf was the highest (30.6%), followed by that with host larvae at 4 dpf (25.7%), 5 dpf (16.1%) and 7 dpf (1.7%) (Figure 2F). Nevertheless, the survival rate of 3 dpf recipients at 3 weeks post transplantation (wpt) was just 36.7%, significantly lower than the survival rates with of SSCT larvae with 4 dpf, 5 dpf or 7 dpf host (Figure 2G). In order to solve the paradox between high fertility rate and low survival rate, we defined a success index of SSCT, which is the product of the percentage values (fertility rate X survival rate), and found that the success index was the highest when using 4 dpf zebrafish as the SSCT host (Figure 2H). Therefore, 4 dpf zebrafish larvae were used as recipients for SSCT.

Therefore, we optimized the cross-subfamily SSCT procedure between *Gr* and zebrafish on the aspects of both donors and hosts. In brief, we injected *cas9-UTRnos3* mRNA that targeted the donor germ cells and *zbuc* mRNA to induce donor germ cells, and utilized 4 dpf zebrafish larvae as the optional recipients.

### 2.3 Generation of functional *Gr* sperm in zebrafish surrogates

To assess whether donor *Gr* SSCs could reconstitute spermatogenesis in sterile zebrafish recipients in a relative short maturation period, we conducted histological analysis of the testes of WT *Gr*, zebrafish, SSCT positive zebrafish as well as *dnd* MO injected zebrafish at 40 dpf, 2 mpf, 3 mpf and 4 mpf to trace the process of spermatogenesis. For WT *Gr*, at 40 dpf, the gonad was naive with plenty of undifferentiated germ cells (Figure 3A, a); at 2 mpf, only a small number of spermatogonia differentiated into primary spermatocytes through meiosis (Figure 3A, b, arrow); and large numbers of spermatozoa could not be observed until 4 mpf (Figure 3A, c-d). As to the zebrafish testis, at 40 dpf, many dark blue spermatocytes have emerged (Figure 3A, e); at 2 mpf, large numbers of spermatids can be seen in zebrafish testis (Figure 3A, f); and at 3 mpf, millions of mature spermatozoa formed in the seminiferous cyst and began to release (Figure 3A, g). All these indicated that the maturation period of *Gr* testis is at least one month later than that of zebrafish testis. However, in SSCT zebrafish testis, the process of spermatogenesis resembled that of zebrafish testis, since the spermatids emerged in the SSCT testes at as early as 2 mpf (Figure 3A, j), and the sperm matured at 3 mpf (Figure 3A, k). In contrast, no spermatogenesis could be observed in non-transplanted control zebrafish testis throughout the maturation period (Figure 3A, m-p). Therefore, by SSCT, we could significantly shorten the spermatogenesis period of *Gr* in zebrafish host.

**Figure 3.**
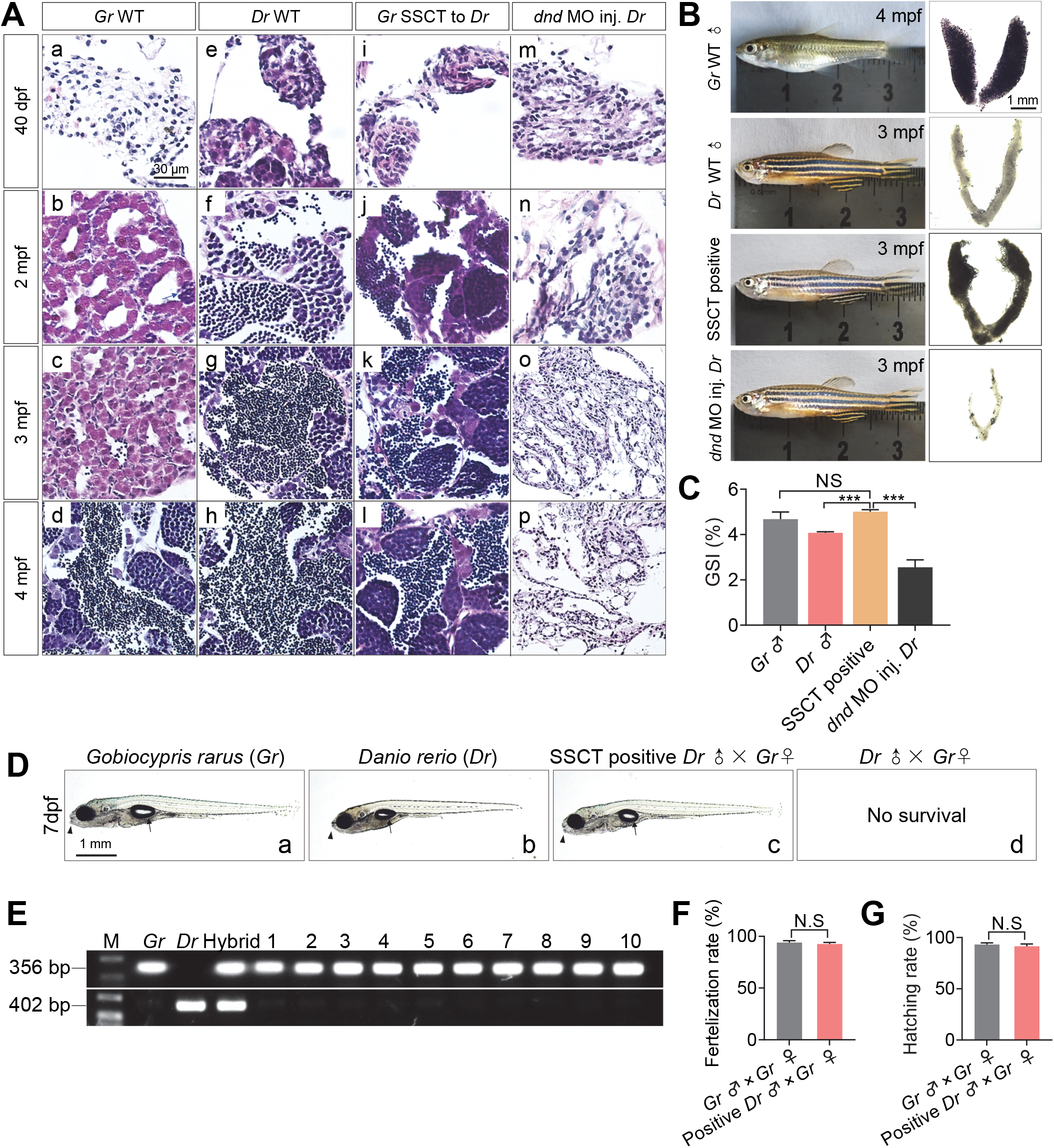
Fast generation of functional *gobiocypris rarus* sperm in zebrafish surrogates. **A**: Process of spermatogenesis of wild type (WT) *Gr*, WT *Dr*, SSCT positive *Dr* and *dnd* MO injected *Dr* testes at 40 dpf, 2 mpf, 3 mpf and 4 mpf. **B**: Morphology of the adult males of WT *Gr*, WT *Dr*, SSCT positive *Dr* and *dnd* MO injected *Dr* and their testes. **C**: The gonadosomatic Index (GSI) of WT *Gr*, WT *Dr*, SSCT positive *Dr* and *dnd* MO injected *Dr*. **D**: Morphology of the larvae crossed by *Gr* male and female (a), *Dr* male and female (b), SSCT positive *Dr* male and *Gr* female (c) at 7 dpf, respectively. The progeny of *Dr* male and *Gr* female cannot even survive to 7 dpf (d). Note that the progeny of SSCT positive *Dr* male and *Gr* female looked exactly as 7 dpf *Gr* larvae in the opened mouth (arrowheads) and the the shape of swim bladder (arrows). **E**: PCR amplification of *chd* genomic DNA fragments in larvae obtained from different parents. The 356 bp and 402 bp amplicons were obtained using specific primers for *Gr* and *Dr chd* respectively. M represents a molecular weight marker; *Gr* represents genome from *Gr* progeny; *Dr* represents genome from zebrafish progeny; Hybrid represents genome from larva crossed by *Dr* male and *Gr* female; 1-10 represent genome from larva crossed by 10 SSCT positive *Dr* males and *Gr* females. **F**: Fertilization rate of the embryos crossed by SSCT positive *Dr* males and *Gr* females was comparative to those of embryos crossed by *Gr* males and *Gr* females. **G**: Hatching rate of the larvae crossed by SSCT positive *Dr* males and *Gr* females was comparative to those of larvae crossed by *Gr* males and *Gr* females. Abbreviations: *Gr, gobiocypris rarus*; *Dr, danio rerio*; SSCT, spermatogonial stem cell transplantation; *dnd, dead end*; MO, morpholino; dpf, days post fertilization.

On the exterior appearance, there was no difference between SSCT positive zebrafish males and WT zebrafish males or the *dnd* MO injected zebrafish males. However, under stereomicroscope, the SSCT positive testis looked non-transparent, which was similar to the testis of WT *Gr*, and quite different from that of the host WT zebrafish testis, which showed to be transparent (Figure 3B). The gonadosomatic index (GSI) of SSCT positive zebrafish was comparative to that of *Gr*, which was significantly higher than that of the *dnd* MO injected zebrafish and even that of the WT zebrafish (Figure 3C).

To identify whether the SSCT positive zebrafish males could produce functional *Gr* sperm, we mated them with *Gr* females by artificial insemination. As a control, the progeny produced by zebrafish sperm and *Gr* eggs could never survive to 7 dpf (Figure 3Dd), while the progeny generated by SSCT positive zebrafish males and *Gr* females could not only reach 7 dpf (Figure 3D), but also develop to adulthood (Figure S3). The larval fish resulting from SSCT positive zebrafish males and *Gr* females looked the same as the *Gr* larvae in morphology (Figure 3Da-c, arrowheads and arrows). PCR analysis of the genomic DNA validated that the larvae crossed by SSCT zebrafish males and *Gr* females were indeed *Gr* larvae, as revealed by amplification of *chd* gene fragments using *Gr* and zebrafish specific primers (Table S1) (Figure 3E). All these demonstrate that the SSCT positive zebrafish males produced functional *Gr* sperm but not zebrafish sperm. Additionally, the fertilization rate (Figure 3F) and hatching rate (Figure 3G) of the embryos crossed by SSCT zebrafish males and *Gr* females were comparative to those resulting from males and females of *Gr*. To conclude, functional *Gr* sperm could be produced in SSCT positive zebrafish recipients and the progress of spermatogenesis was accelerated in zebrafish recipients.

### 2.4 Colonization, proliferation, and differentiation of *Gr* SSCs in zebrafish recipients

In order to better understand the developmental dynamics of the transplanted SSCs in zebrafish recipients, we analyzed the colonization, proliferation, and differentiation *Gr* germ cells by different methods. Firstly, at 3 wpt, by using red fluorescent labeling of donor cells, we observed that the fluorescent donor cells colonized in recipient gonadal region below the swimming bladder (Figure 4A, a), while no fluorescence was observed in nontreated control larvae (Figure 4A, b). Further dissection and observation revealed that numerous fluorescent donor cells colonized in the host zebrafish gonad (Figure 4A, a’), while no fluorescence was observed in non-transplanted control gonad (Figure 4A, b’). The colonization rate varies from 25% to 85% according to different donors and recipients (Table 1). To study the proliferation of donor SSCs, we conducted EdU assay and found that numerous proliferative VASA positive germ cells existed in SSCs colonized zebrafish gonads (Figure 4B, b), just mimicking those in WT zebrafish gonads (Figure 4B, a), whereas there was no proliferative germ cell in non-transplanted control gonads (Figure 4B, c).

**Table 1.**
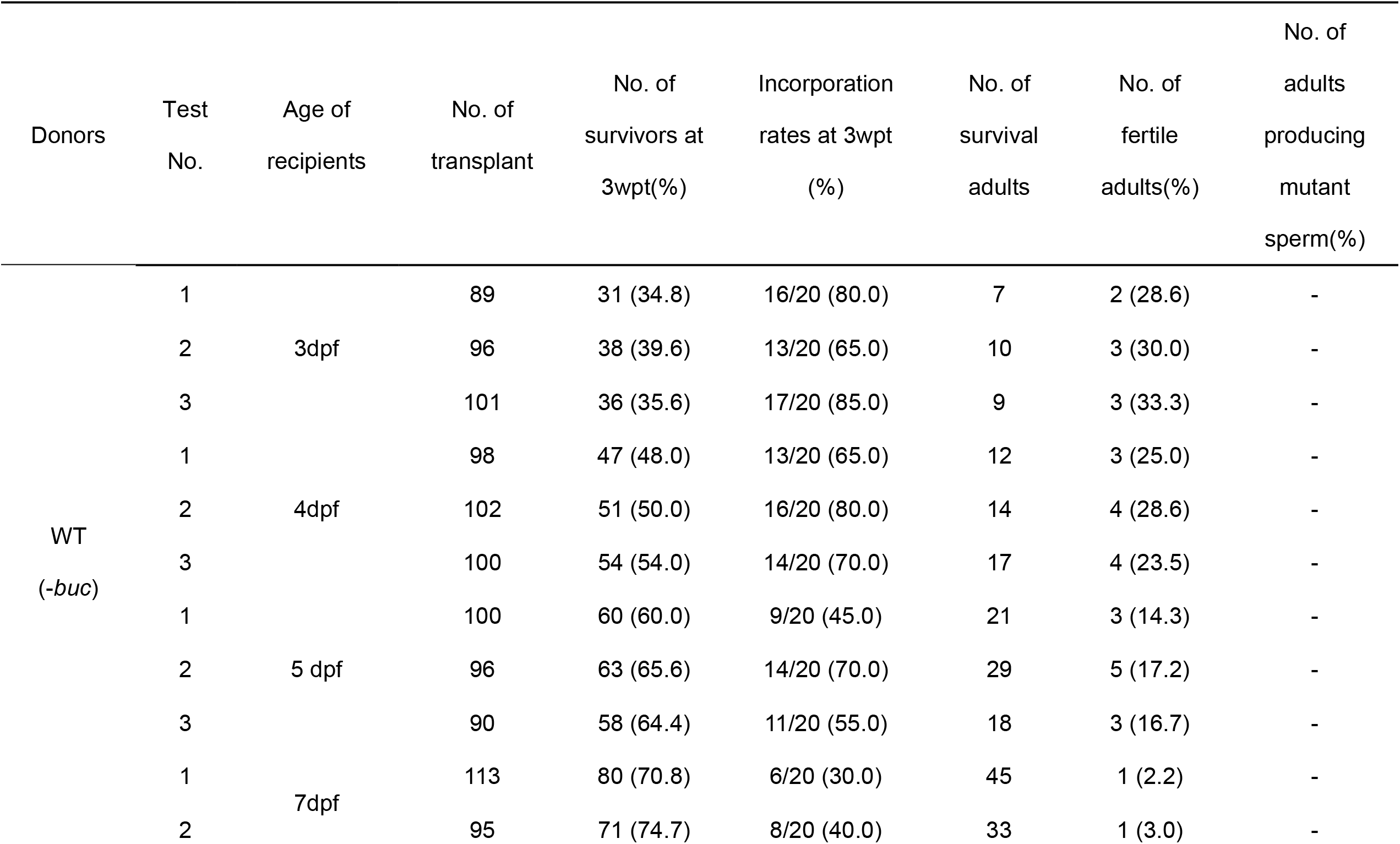
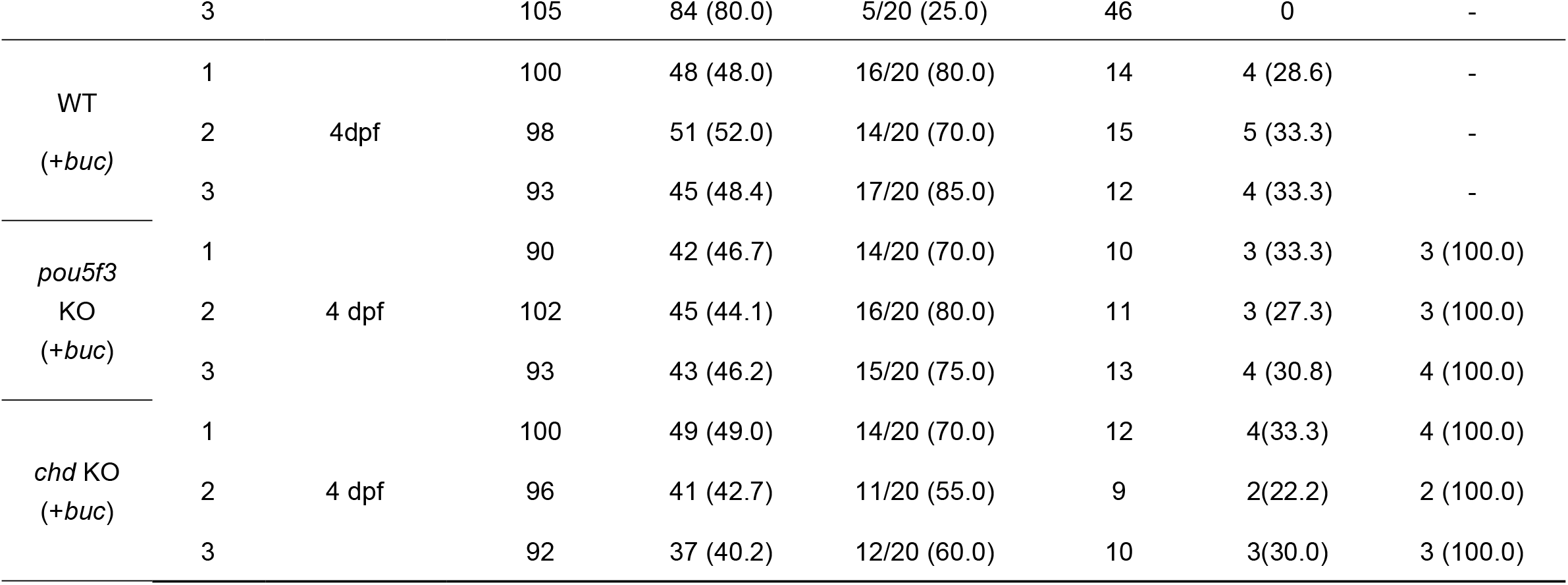
Statistics of survival rate, incorporation rate and the fertility rate of grown adults after SSCT

**Figure 4.**
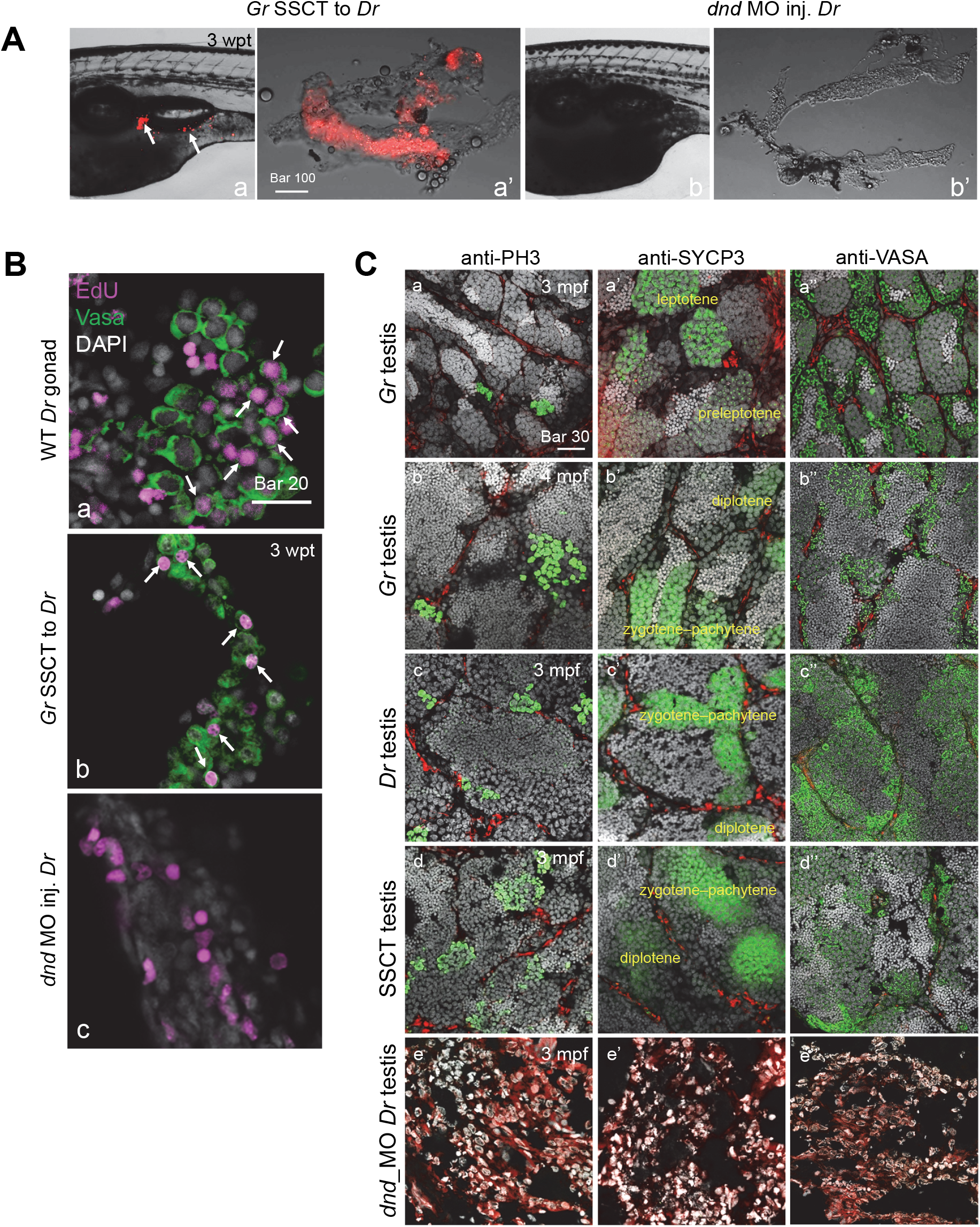
Colonization, proliferation and differentiation of *gobiocypris rarus* spermatogonia in zebrafish recipients. **A**: Colonization of red fluorescent labeled donor cells into recipient genital ridge (a, arrows) and gonad (a’) at 3 wpt, while no red fluorescent labeled donor cells were observed neither in non-transplanted *dnd* MO injected recipient zebrafish (b) nor its gonad (b’). Note that the non-transplanted recipient zebrafish gonad was lamellar, while SSCs transplanted zebrafish gonad looks plump. **B**: Proliferative VASA-positive germ cells were observed both in wild type (WT) (a) and SSCs transplanted (b) zebrafish gonads but not in non-transplanted recipient zebrafish gonad injected with *dnd* MO (c). Arrows showed the proliferative VASA-positive germ cells. **C**: Immunofluorescence detection with PH3 and SYCP3 antibodies showed that as in WT *Gr* and *Dr* testis, mitosis and meiosis were normal in SSCT positive *Dr* testis. Signals of these two antibodies were not detectable in *dnd* MO injected *Dr* testis. As the same with *Gr* and *Dr* testis, VASA expressed in all stages of germ cells except for spermatozoa and spermatids in SSCT positive *Dr* testis. *dnd* MO injected *Dr* testis was used as a negative control. Note that 3 mpf *Gr* testis was immature as most of the spermatocytes were at the preleptotene or leptotene stage (b), indicating their beginning of miosis initiation, while the spermatocytes of 4 mpf *Gr* testis, 3 mpf *Dr* testis and SSCT positive testis were at the zygotene–pachytene or diplotene stage. Additionally, 3 mpf *Gr* testis contained most of the spermatogonia and spermatocytes with just a few spermatids, while numerous spermatids and spermatozoa could be observed in 4 mpf *Gr* testis, 3 mpf *Dr* testis and SSCT positive testis. Abbreviations: wpt, weeks post transplantation; *dnd, dead end*; MO, morpholino; SSCs, spermatogonia stem cells; *Gr, gobiocypris rarus*; *Dr, danio rerio*; mpf, months post fertilization.

To further characterize the proliferation and differentiation of transplanted germ cells in SSCT fish, we conducted immunofluorescent staining of the testes with a mitosis marker PH3 (Perez-Cadahia et al., 2009), which indicates cell proliferation, and a miosis marker SYCP3 (Yuan et al., 2000), which indicates differentiation of germ cells. In the developing testis, a lot of cells undergoing mitosis and miosis in the SSCT positive testis (Figure 4C, d, d’), mimicking what was observed in *Gr* testis (Figure 4C, a-b, a’-b’) and zebrafish adult testis (Figure 4C, c, c’). Immunostaining of VASA confirmed that the transplanted *Gr* SSCs resumed spermatogenesis of infertile zebrafish males, and various germ cells including spermatogonia, spermatocytes, spermatids, and spermatozoa were observed in the SSCT positive testis (Figure 4C, d’’), consistent with what was seen in WT *Gr* (Figure 4D, b’’) and zebrafish testis (Figure 4D, c’’). In contrast, at 3 mpf, the *Gr* testis only contained large numbers of spermatogonia and spermatocytes with a few spermatids (Figure 4D, a, a’, a’’), and large amounts of spermatocytes at the preleptotene or leptotene stage, indicating the initiation of meiosis (Figure 4D, a’). These demonstrate that the *Gr*-originated germ cells could efficiently proliferate and differentiate in the zebrafish host testes, and the maturation period of SSCT testis resembled that of zebrafish testis but not *Gr* testis.

### 2.5 The surrogate testis consists of *Gr*-derived germ cells and zebrafish-derived gonadal somatic cells

To further characterize the composition of different gonadal cells in the surrogate testis, we performed RNA in situ hybridization analysis against species-specific markers of germ cells (*vasa*), and two types of gonadal somatic cells, the Leydig cells (*insl3*) (Ivell and Bathgate, 2002) and Sertoli cells (*gsdf*) (Sawatari et al., 2007). Firstly, we showed the effectiveness and specificity of the species-specific markers. As shown in Figure 5, *Gr vasa* and zebrafish *vasa* could only label the germ cells of *Gr* testis and zebrafish testis (Figure 5A, a-b’), respectively; *Gr insl3* and zebrafish *insl3* only labeled the Leydig cells of *Gr* testis and zebrafish testis (Figure 5B, a-b’), respectively; and *Gr gsdf* and zebrafish *gsdf* could only label the Sertoli cells of *Gr* testis and zebrafish testis (Figure 5C, a-b’), respectively. Interestingly, in the SSCT testis, only *Gr* germ cells but not zebrafish germ cells could be detected (Figure 5A, c-c’); while both the Leydig cells and Sertoli cells were from the host zebrafish (Figure 5B, c-c’, C, c-c’), suggesting that the SSCT testis was a reconstructed gonad, consisting of *Gr*-derived germ cells and zebrafish-derived gonadal somatic cells. In contrast, in the *dnd* morphant host testis, there was no labeling signal of germ cells (Figure 5A, d-d’), and completely disrupted structures could be found with disorganized expression of zebrafish-specific *insl3* and *gsdf* (Figure 5B, d-d’, C, d-d’). RT-PCR amplification of partial CDS of *vasa* gene from cDNA samples using *Gr* and zebrafish *vasa* specific primers (Table S1) further validated that the germ cells in SSCT positive testis were derived from transplanted *Gr* SSCs but not endogenous zebrafish PGCs (Figure S4).

**Figure 5.**
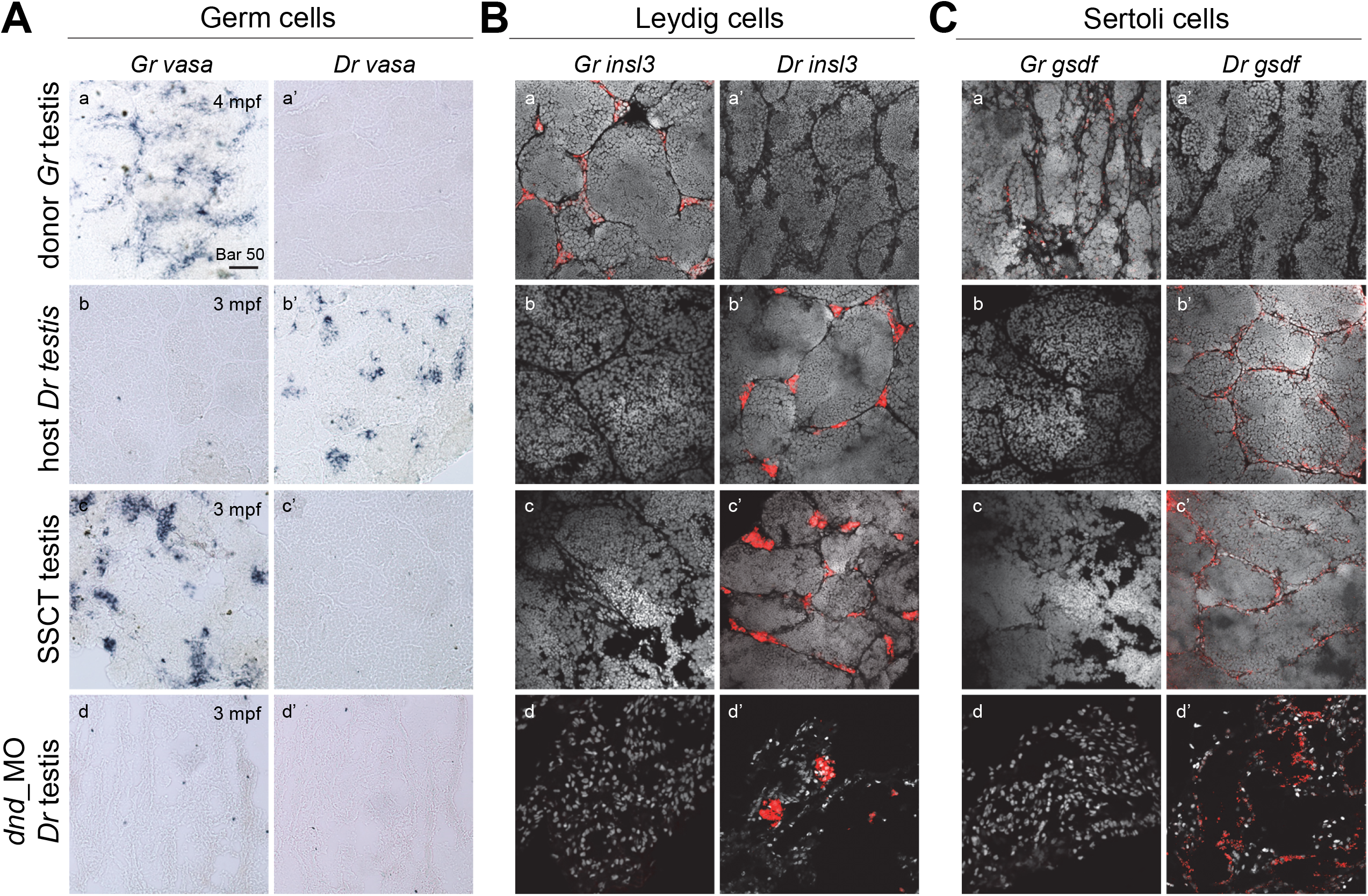
SSCT positive testis consisted of donor-derived germ cells and recipient-derived gonadal somatic cells. **A**: In situ hybridization on sections using *Gr* and *Dr* specific *vasa* probe showed that they specifically expressed in *Gr* and *Dr* testes respectively, while *Gr vasa* but not *Dr vasa* expressed in SSCT positive *Dr* testis. Neither *Gr vasa* nor *Dr vasa* were detected in *dnd* MO injected *Dr* testis. **B**: FISH on sections using *Gr* and *Dr* specific *insl3* probe showed that they specifically expressed in *Gr* and *Dr* testes respectively. *Dr insl3* was also detected in SSCT testis and *dnd* MO injected *Dr* testis. **C**: FISH on sections using *Gr* and *Dr* specific *gsdf* probe showed that they specifically expressed in *Gr* and *Dr* testes respectively. *Dr gsdf* was also detected in SSCT testis and *dnd* MO injected *Dr* testis. Abbreviations: SSCT, spermatogonial stem cell transplantation; *Gr, gobiocypris rarus*; *Dr, danio rerio*; *dnd, dead end*; MO, morpholino; FISH, fluorescence in situ hybridization.

Taken together, we revealed that the *Gr* SSCs could not only colonize and proliferate in the zebrafish recipients, but also support the development and function of zebrafish gonadal somatic cells, and finally undergo normal spermatogenesis and differentiate into spermatozoa in the reconstructed functional testes.

### 2.6 Morphology and swimming behavior of sperm derived from SSCT resembles that of hosts but not donors

Although the SSCT positive zebrafish could produce functional *Gr* sperm, we were interested in whether there was any difference between the sperm derived from the surrogate zebrafish and the *Gr* males. By scanning electron microscopy (SEM) analysis, the morphology of SSCT sperm looked more similar to zebrafish sperm than *Gr* sperm, as the first two had the structure of protruded mitochondrial sheath (Figure 6A, red boxes) and they owned elliptical ball-shaped heads while *Gr* sperm head looked much more round with no typical mitochondrial sheath (Figure 6A,a’, b’, c’). Moreover, the average head diameter of SSCT sperm (1.55 μm) was significantly larger than that of *Gr* sperm (1.46 μm), and much closer to that of zebrafish sperm (1.65 μm) (Figure 6B). The average tail length of SSCT sperm (30.32 μm) was almost similar to that of zebrafish sperm (30.80 μm), but significantly shorter than that of *Gr* sperm (32.42 μm) (Figure 6C). The center of their flagella was the axial filament with a typical “9×2+2” microtubule structure (Figure S5A, a, b, c), and the tail diameter showed no significant difference among *Gr*, zebrafish and SSCT sperm (Figure S4B). Transmission electron microscopy (TEM) analysis revealed that the membrane of head showed serrated phenotype both in *Gr* (Figure S5A, a’) and SSCT sperm (Figure S5A, c’) while it was smooth in zebrafish sperm (Figure S5A, b’). Zebrafish sperm possessed overt cytoplasm in the front of the head, which was not seen in the *Gr* head nor the SSCT sperm head. Statistics showed that the mitochondria number was a little bit lower in *Gr* and SSCT sperm, when compared to that of zebrafish sperm (Figure S5C).

**Figure 6.**
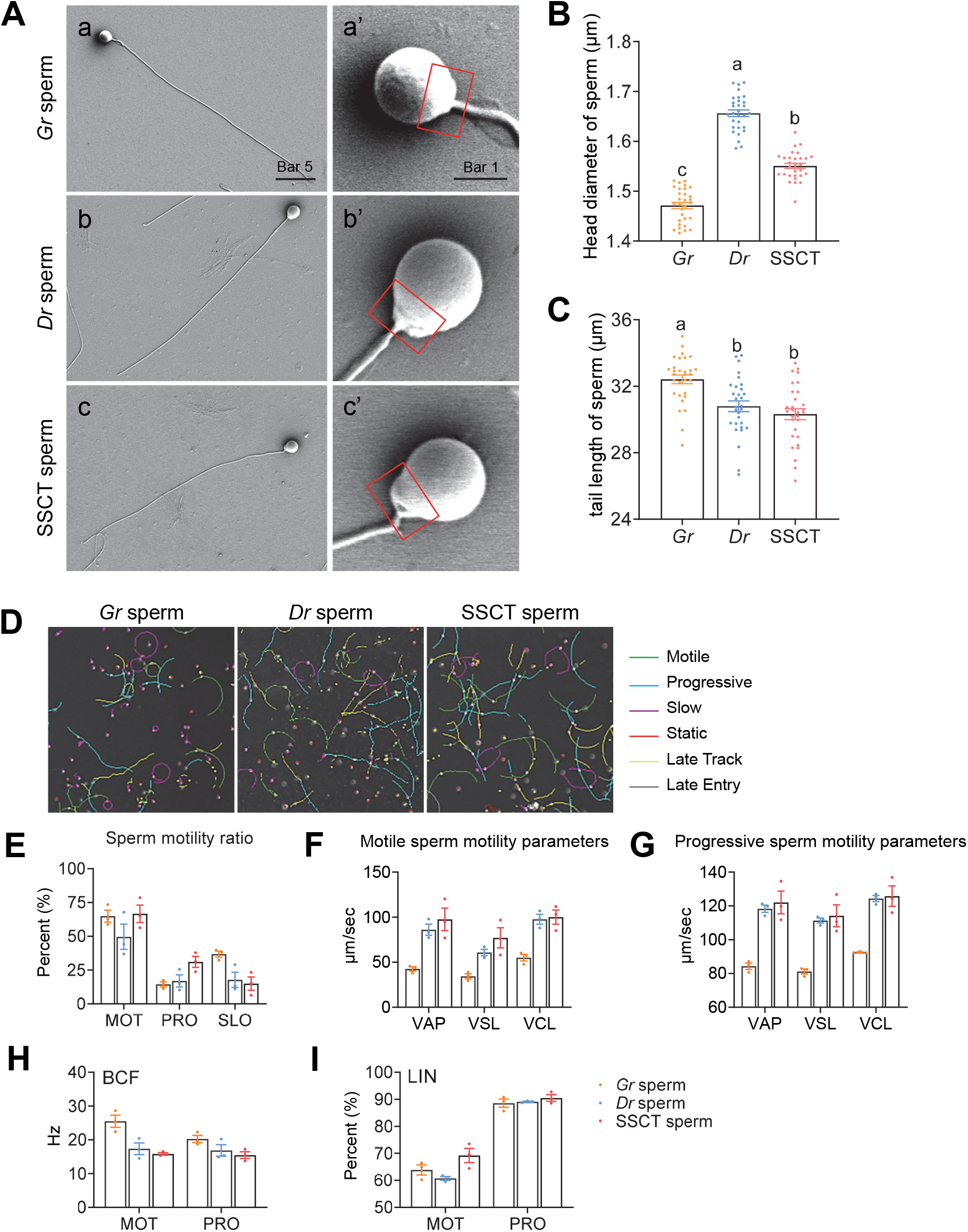
Morphology and activity of sperm derived from SSCT positive zebrafish resembles that of hosts but not donors. **A**: SEM images of sperms derived from wild type (WT) *Gr*, WT *Dr* and SSCT positive *Dr* testes. a’, b’ c’ were magnifications of the sperm heads in a, b, c, respectively. Red frames showed the mitochondrial sheath of different sperms. **B**: Head diameter of sperms derived from WT *Gr*, WT *Dr* and SSCT positive *Dr* testes. **C**: Tail length of sperms derived from WT *Gr*, WT *Dr* and SSCT positive *Dr* testes. **D**: Different kinestates of *Gr, Dr* and SSCT sperms. **E**: Percentage of motile (MOT), progressive (PRO) and slow (SLO) sperms of *Gr, Dr* and SSCT sperms. **F**: Parameters of motile sperm motility of *Gr, Dr* and SSCT sperms, including VAP (average path velocity), VSL (straight line velocity) and VCL (curvilinear velocity). **G**: Parameters of progressive sperm motility of *Gr, Dr* and SSCT sperms, including VAP, VSL and VCL. **H**: Beating cross frequency (BCF) of the motile and progressive sperms of *Gr, Dr* and SSCT sperms. **I**: Linearity (LIN) of the motile and progressive sperms of *Gr, Dr* and SSCT sperms. Abbreviations: SSCT, spermatogonial stem cell transplantation; SEM, scanning electron microscopy; *Gr, gobiocypris rarus*; *Dr, danio rerio*.

Subsequently, the activity of *Gr*, zebrafish and SSCT sperm was measured and the trajectory map of SSCT sperm resembled that of zebrafish sperm but not *Gr* sperm (Figure 6D). Statistics showed that the ratio of motile SSCT sperm was comparative to that of *Gr* sperm and higher than that of zebrafish sperm, but the ratio of progressive SSCT sperm was higher than that of both *Gr* and zebrafish sperm while the ratio of slow SSCT sperm was comparative to that of zebrafish sperm and lower than that of *Gr* sperm (Figure 6E). Notably, the motility parameters like average path velocity (VAP), straight line velocity (VSL) and curvilinear velocity (VCL) of both motile (Figure 6F) and progressive (Figure 6G) SSCT sperm were comparative to those of zebrafish but not *Gr* sperm. Furthermore, the beat cross frequency (BCF) of both motile and progressive SSCT sperm was like that of zebrafish sperm but lower than that of *Gr* sperm (Figure 6H). However, the linearity (LIN, VSL/VCL) of motile SSCT sperm was higher than both that of *Gr* and zebrafish sperm while the LIN of progressive sperm of them three was similar (Fig.6I). To sum up, the morphology and activity of *Gr* sperm produced by SSCT positive zebrafish recipients resembles that of zebrafish hosts but not *Gr* donors.

### 2.7 Efficient production of *pou5f3* and *chd* mutated *Gr* sperm in zebrafish surrogates

In our previous study, we have successfully generated donor-derived genome edited gametes through combination of CRISPR/Cas9 gene knockout technology and allogenic PGCT in zebrafish (Zhang et al., 2020). We further inquired whether functional genome edited *Gr* sperm could be achieved by cross-subfamily SSCT into zebrafish surrogates. Here we chose two genes, *pou5f3* (Burgess et al., 2002) and *chd* (Hammerschmidt et al., 1996), which are essential for brain and dorsal development, respectively. The genome edited *Gr* testicular cells were dissociated, and the SSCs were purified and used for xenogeneic SSCT into zebrafish surrogates. When the transplanted zebrafish grew to adulthood at 3 mpf, SSCT positive individuals were screened out by mating them with WT zebrafish females one by one. To evaluate the mutation efficiencies of sperm produced by SSCT positive males, the target sequences of each embryo crossed by a SSCT positive male and a zebrafish female were analyzed.

When the *pou5f3* knocked out *Gr* SSCs were used as donors for SSCT, we finally obtained 10 zebrafish surrogates producing *Gr* sperm and all of them could produce *pou5f3* mutated *Gr* sperm. Surprisingly, the mutation efficiencies of the SSCT sperm reached as high as 100% in all the 10 surrogates (Figure 7A), and the mutation types were shown in Table S2. Furthermore, 51.8% of the progeny showed a typical phenotype with shortened trunk and aberrant tail (Figure 7B, b) when crossing #1 SSCT positive zebrafish male with a *Gr pou5f3* mutated heterozygote female (−4 bp). Sanger sequencing showed that C1 embryos with WT like phenotype were all heterozygotes while C2 embryos contained only mutant alleles for the target sequences (Table 2). Phenotypic analysis showed that C1 embryos displayed normal brain structures including tectum (t), cerebellum (c), hindbrain (h) and otic placode(o) at 30 hpf (Figure 7B, a’), while in C2 embryos, the hindbrain was disorganized (Figure 7B, b’, red arrowhead) and the otic placode was reduced in size containing only one otolith (Figure 7B, b’, blue arrowhead). α-tubulin staining further validated the disorganization of brain structure in C2 embryos at 2 dpf (Figure 7C). Detection of the expression of some relevant molecular makers by WISH showed that *pou5f3* barely expressed in mutant embryos neither at shield nor bud stage (Figure 7D, left two panels), while the expression of *krox20*, a marker for midbrain and hindbrain, and *wnt1*, a marker for midbrain-hindbrain boundary (MHB), was reduced markedly in *pou5f3* mutated embryos (Figure 7D, right two panels). These results were consistent with those found in zebrafish *pou5f3* mutants (Burgess et al., 2002), suggesting that zygotic *pou5f3* is also essential for the brain development of *Gr*.

**Table 2.**
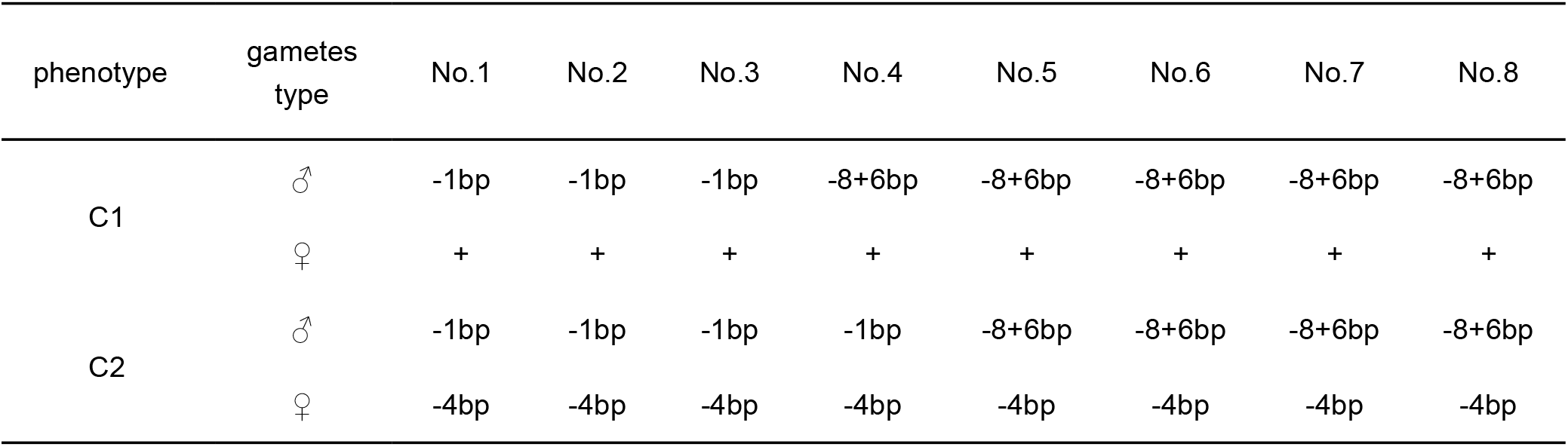
Genotyping of the F1 embryos crossed by 1# SSCT positive male producing *pou5f3* mutated *Gr* sperm and a *Gr pou5f3* mutant heterozygous female (−4bp)

**Figure 7.**
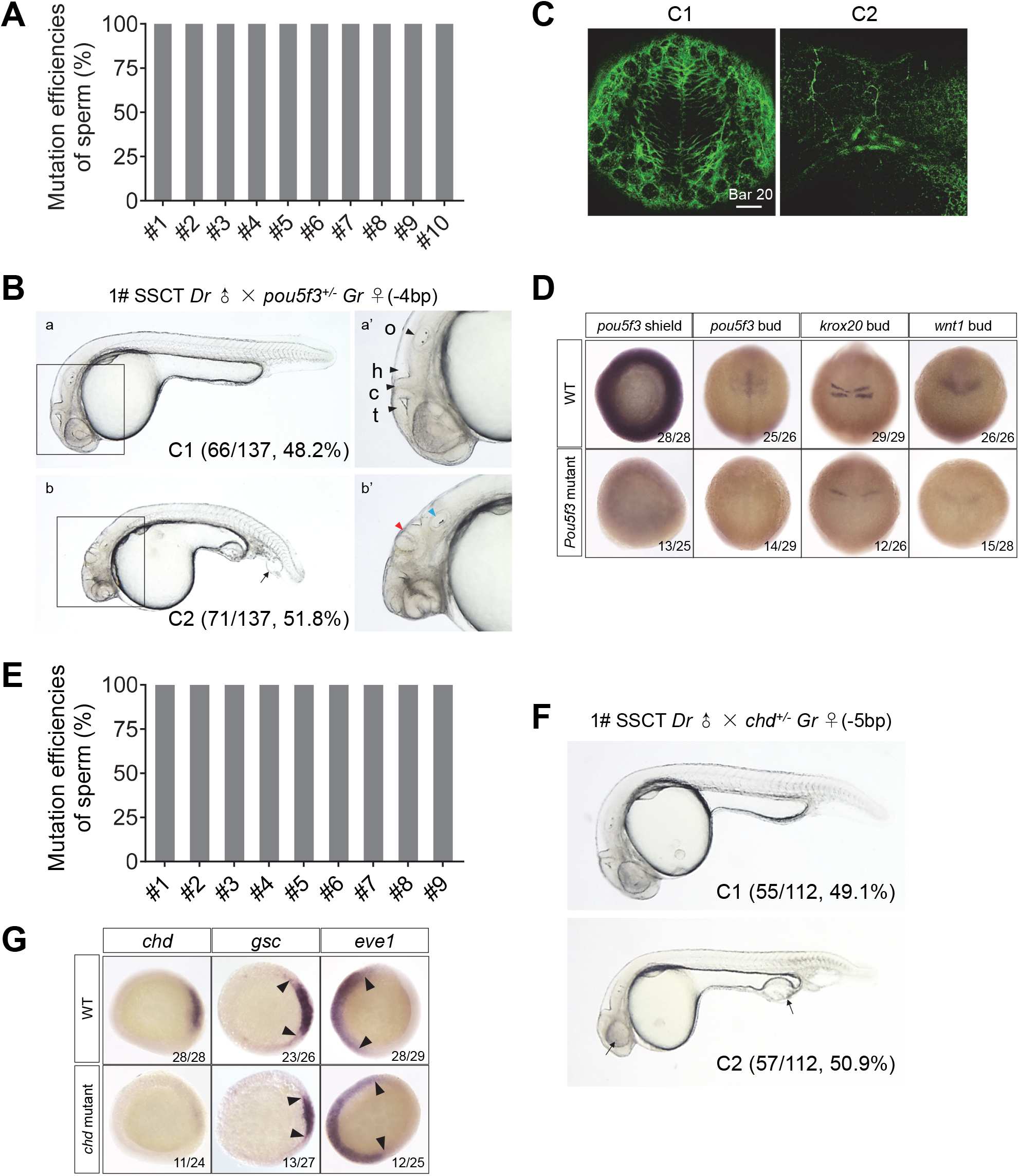
Efficient production of *pou5f3* and *chd* mutated *Gr* sperm in zebrafish surrogates. **A**: Mutation efficiencies of sperms from 10 SSCT positive zebrafish adult males producing *Gr pou5f3* mutated sperms. **B**: The phenotypes of F1 offspring crossed by #1 SSCT *Dr* male producing *Gr pou5f3* mutated sperms and a *Gr pou5f3* mutated heterozygote female (−4 bp). C1 shows the wild type (WT) like phenotype (a), C2 shows altered morphology of the tail (arrowhead) and shortened trunk at 30 hpf (b). Note that C1 embryos display normal brain structures including tectum (t), cerebellum (c), hindbrain (h) and otic placode(o) at 30 hpf (a’). The hindbrain displays disorganization (red arrowhead) and the otic placode is reduced in size, containing only one otolith (blue arrowhead) in embryos with C2 phenotype (b’), indicating that *pou5f3* functions in *Gr* brain development. **C**: α-tubulin staining recognizes the well-organized axonal scaffold in the hindbrain of C1 embryos at 2 dpf while *pou5f3* mutant embryos with C2 phenotype show strong disorganization of the axonal scaffold within the hindbrain. **D**: *pou5f3* expresses all over the germ ring at shield stage and specifically expresses in the neuroectoderm at bud stage in wild type (WT) embryos, but in mutant embryos no expression of *pou5f3* was detected at these two stages. *krox20* was not properly initiated in rhombomere (r) 3 and r5 and failed to fuse at the midline in mutant embryos. Expression of a marker of the MHB *wnt1 was strongly reduced in pou5f3* mutant embryos compared to WT embryos. **E**: Mutation efficiencies of sperms from 9 SSCT positive zebrafish adult males producing *Gr chd* mutated sperms. **F**: The phenotypes of F1 offspring crossed by #1 SSCT *Dr* male producing *Gr chd* mutated sperms and a *Gr chd* mutated heterozygote female (−5 bp). C1 showed the WT like phenotype, C2 showed smaller eyes and enlarged blood island, a typical phenotype of zygotic mutant. **G**: *chd* was barely detected at the organizer in *chd* mutant embryos compared to its strong expression in WT embryos. Expression of the marker of dorsal region *gsc* was reduced while expression of the marker of ventral region *eve1* was enhanced in mutant embryos.Abbreviations: SSCT, spermatogonial stem cell transplantation; *Gr, gobiocypris rarus*; *Dr, danio rerio*; MHB, midbrain-hindbrain boundary.

Chd has been known as a major antagonist of BMP signaling in early development and is essential for dorsal development (Nasevicius and Ekker, 2000; SchulteMerker et al., 1997). Similarly, when *chd* knocked out *Gr* SSCs were used as donors for SSCT, 9 zebrafish surrogates produced *chd* mutated *Gr* sperm with 100% efficiencies (Figure 7E). The #1 SSCT positive zebrafish male was crossed with a *Gr chd* mutated heterozygote female (−5 bp) and 50.9% of the C2 progeny showed a typical phenotype of moderate ventralization (Figure 7F), which was consistent with that of zebrafish *chd* mutants (Zhang et al., 2016). Sanger sequencing further proved that the C2 embryos only contained mutant alleles (Table 3). WISH results revealed that *chd* was scarcely expressed in the mutant embryos and the expression of dorsal marker *gsc* was reduced while the maker for ventral development *eve1* extended dorsally (Figure 7G). Therefore, the anti-BMP function of Chd is conserved between zebrafish and *Dr*. Taken together, these validated that *Gr* mutant sperm could be obtained efficiently in zebrafish host within a relative short maturation period, by CRISPR/Cas9 targeted SSCT technology.

**Table 3.**
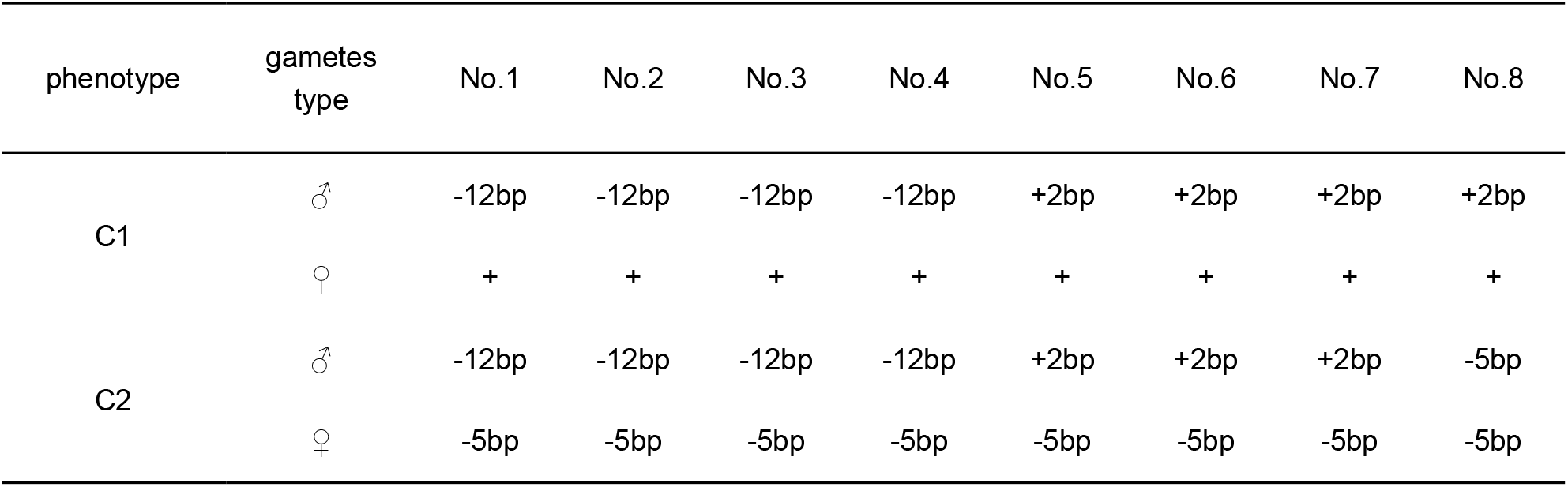
Genotyping of the F1 embryos crossed by 1# SSCT positive male producing *chd* mutated *Gr* sperm and a *Gr chd* mutant heterozygous female (−5bp)

## 3. Discussion

It is generally proposed that combining the recent biotechnological innovations, such as genome editing and surrogate broodstock technologies, could be utilized to expedite aquaculture breeding in the future (Gratacap et al., 2019; Houston et al., 2020; Sun and Zhu, 2019). Recently, by combining the CRISPR/Cas9 gene knockout and PGCs transplantation technologies, we successfully generated zebrafish maternal-zygotic mutants of embryonic lethal genes at F1 generation (Zhang et al., 2020). In the current study, by utilizing two small cyprinid fish from different subfamilies, we successfully generated genome edited sperm of *Gr* in zebrafish host by SSCT. This is the first report on production of genome-edited gametes via surrogate broodstock technology between two fish species, which provides a paradigm for future genetic breeding of aquaculture species, with those innovative biotechnologies, such as genome editing and surrogate technology.

In previous studies of fish surrogate reproduction, different sources of donor cells have been utilized for germline transplantation, such as PGCs-containing blastoderm cells, single PGC, SSCs, or OSCs, and different methods of transplantation have been developed, such as peritoneal cavity transplantation into fish larvae and genital pore transplantation into fish adults (Goto and Saito, 2019). It has been shown that the transplantation between different individuals or different species could be successful when using different combinations of germline donor and transplantation method, and the success rate was usually negatively correlated with the distance of phylogenetic relationship between donor species and host species. Among all the combinations, SSCT into peritoneal cavity of fish larvae is considered as the optional strategy of surrogate reproduction, since the SSCs could be easily isolated, cultured, cryopreserved and even expanded after cryopreservation (Iwasaki-Takahashi et al., 2020). However, when the donor and host species were from different subfamilies or two species with longer phylogenetic distance, SSCT has never been succeeded. In the present study, although zebrafish and *Gr* are from different subfamilies, and even the breeding time differs between these two fish species (*Gr* usually spawn at night while zebrafish lay eggs in the morning), we show that *Gr* SSCs could successfully colonize, proliferate, and differentiate in host zebrafish, which finally leads to the efficient production of functional sperm of *Gr* in zebrafish host. Through a series of optimization steps, such as germ cell induction, SSCs isolation, and comprehensive evaluation of the success index of SSCT, we significantly optimized the success rate of SSCT between *Gr* and zebrafish. Therefore, our study has further extended the success of SSCT to two phylogenetically distant species, which belong to different subfamilies.

The progress of colonization, proliferation, and differentiation of GSCs in the surrogate host is critical for the final success of surrogate reproduction, thus each step of this progress should be clearly described and characterized. However, in previous studies of fish surrogate reproduction, this biological process has rarely been described and studied. In our study, by using a combination of histology (Figure 3A), immunochemistry (Figure 4B and C), and RNA in situ hybridization (Figure 5) techniques, we extensively investigated the development of the surrogate gonads, and compared it with that of the host zebrafish and donor *Gr*. Our study showed that the developmental timing and overall morphology of the surrogate testes resembled those of the host zebrafish, instead of *Gr*, therefore the surrogate zebrafish could produce mature *Gr* sperm in 3 months, shorter than the maturation time of *Gr*, which is 4 months. Furthermore, the colonization and proliferation of *Gr* SSCs in host testes, and the spermatogenesis of the surrogate testes were clearly demonstrated. More interestingly, our study explicitly showed that the surrogate testes were composed of the germ cells derived from *Gr* and testicular somatic cells (Leydig cells and Sertoli cells) of zebrafish, which further revealed the conservation of signaling pathway controlling testis development and spermatogenesis in different fish species. All these studies undoubtedly demonstrated the success of surrogate production of *Gr* sperm in a short maturation period, by SSCT into zebrafish host.

In the field of surrogate reproduction technology, the main obstacle restricting its wide-range application should be the low success rate of GSCT. In order to get the highest success rate for SSCT, we optimized the technique on both aspects of the donor and the host. On one hand, we found that 4 dpf larvae were the most suitable for the surrogate host, as a relatively high fertility rate and an acceptable death rate could be obtained simultaneously after SSCT. This solved the paradox between survival and transplantation efficiency encountered before (Li et al., 2017). On the other hand, *zbuc* mRNA was injected into donor embryos to induce additional PGCs thus elevated the success rate of xenogeneic SSCT to 31.7% when 4 dpf zebrafish larvae were served as recipients (Figure 2E). This data was higher than the success rate obtained in zebrafish allogenic ovarian germ cells transplantation (OGCT) using 2-wk-old sterile hybrid larvae as recipients (18%) (Wong et al., 2011), and even slightly higher than that achieved in zebrafish allogenic SSCT when using the busulfan treated adults as recipients (30%) (Nobrega et al., 2010), or the 9-10 dpf *dead end*-knockout larvae as recipients (5%) (Li et al., 2017). In the future, in order to further elevate the success rate of SSCT technology, in vitro-expanded SSCs might be considered as donors for xenogeneic SSCT (Iwasaki-Takahashi et al., 2020).

The surrogate sperm could only fertilize the eggs from donor species, *Gr*, but not recipient species, zebrafish, demonstrating the complete success of SSCT between these two species. However, on the aspect of morphology and ethological characteristics, the surrogate sperm were markedly different from the *Gr* sperm, and somehow resembled the zebrafish sperm. In previous studies, the morphological characteristics of donor derived eggs from surrogate fish were also different from those of control donor eggs, but resembled those of recipient eggs (Yoshikawa et al., 2018; Yoshikawa et al., 2020). These findings indicate that the gametogenesis will be significantly remolded within the gonadal microenvironment of recipient gonadal somatic cells, resulting in not only the synchronous development of gametes but also their similar morphogenesis with those of the surrogate host. In the future, further study is needed to clarify whether there is any exchange of genetic materials between donor germ cells and host gonadal somatic cells.

By far, a big challenge in the field of fish surrogate reproduction is how to efficiently obtain donor-derived functional eggs. During the past decades, surrogate production of donor derived eggs has been only achieved in a few cases, such as allogenic GSCT in zebrafish (Zhang et al., 2020), rainbow trout (Iwasaki-Takahashi et al., 2020), medaka (Li et al., 2016), Chinese rosy bitterling (Octavera and Yoshizaki, 2019), Yellowtail (Morita et al., 2012) and nibe croaker (Yoshikawa et al., 2017), and xenogeneic GSCT between salmon and trout (Okutsu et al., 2007; Takeuchi et al., 2004). In our study, we successfully obtained functional *Gr* sperm but not eggs with zebrafish recipients. Therefore, it is likely more difficult to obtain donor derived eggs than sperm as the phylogenetic distance between donors and hosts increasing, which may be attributed to that fish oogenesis requires maternally supplied proteins such as vitellogenin and egg envelop proteins synthesized by recipient liver (Lubzens et al., 2010). In the future, in order to efficiently obtain donor derived eggs by surrogate reproduction, we may need to modify the host genome to express vitellogenin and egg envelop proteins from the donor species, and to generate a female-biased population as surrogate host as well.

## 4. Materials and Methods

### 4.1 Ethics

This study was carried out in accordance with Guide for the Care and Use of Laboratory Animals in University of Chinese Academy of Sciences and Institute of Hydrobiology, Chinese Academy of Sciences.

### 4.2 Fish and embryos

The experimental fish used in this study were zebrafish of AB line and WT *gobiocypris rarus*, housed in the China Zebrafish Resource Center, National Aquatic Biological Resource Center (CZRC/NABRC, Wuhan, China) and raised at 28 °C with a 14h:10h light and dark cycle. The embryos for microinjection were harvested from natural fertilization and artificial insemination for zebrafish and *gobiocypris rarus* respectively. The stages of embryonic development refer to the paper (Kimmel et al., 1995).

### 4.3 Target site gRNA design and synthesis

First, we got partial CDS sequences of *gobiocypris rarus pou5f3* and *chd* genes by amplifying from their cDNA using pairs of primers (Table S1) designed according to the conserved regions of several teleosts’ sequences information (https://www.ncbi.nlm.nih.gov/gene/). Then we further confirmed partial genome sequences of these two genes by amplifying from its genome using same pairs of primers. Finally, we designed two target sites for each gene on the website of http://zifit.partners.org/ZiFiT/. By pre-experiment, we eventually got effective target sites:

*pou5f3*-target: GGGACGTCATCCGTGCCCAC<u>CGG</u>.

*chd*-target: GGAGCGCGGCGGAGGCCGGG<u>CGG</u>;

Sequences underlined show PAM (Protospacer adjacent motif). gRNA templates were prepared by PCR (94 °C 5 min; 94 °C 30 s, 60 °C 30 s, and 72 °C 30s,30 cycles; 72 °C 8 min) with gene specific primers (*pou5f3*-gRNA-F2, *chd*-gRNA-F4) and a universal reverse primer gRNA-RP (Table S1) using plasmid pT7-gRNA as template (Chang et al., 2013). After purification, they were transcribed with MAXIscript T7Kit (Ambion, USA).

### 4.4 Microinjection of mRNA and gRNA

The mRNA of *Cas9-UTRnos3, buc-SV40*, and *gfp-UTRnanos3* was prepared as described in our previous study (Zhang et al., 2020). *Cas9-UTRnanos3* mRNA and gene specific gRNAs were injected with dosages of 300 pg and 80 pg per embryo into donor embryos of *Gr*. Zebrafish embryos were injected with 100 μM of *dead end* (*dnd*) antisense morpholino oligonucleotide (5’-GCTGGGCATCCATGTCTCCGACCAT-3’) to eliminate endogenous PGCs according to previous report (Weidinger et al., 2003; Zhang et al., 2020).

### 4.5 Purification and transplantation of spermatogonia

Three testes from 4-months-old *pou5f3* and *chd* KO as well as WT *Gr* adults were minced with scissors and dissociated in 0.25% trypsin (Invitrogen) in 1mL L-15 medium (Sigma-Aldrich) with 0.26 U Librase (Roche), 0.05% Dnase I (Roche) and 10% FBS (BioInd) for 2-00203 hrs at room temperature (RT). The resulting testicular cell suspension was filtered through a 40-μm mesh to remove large debris, washed with PBS, and the cell pellet was resuspended in 1 mL of PBS. The cell suspension was loaded onto a discontinuous Percoll (GE Healthcare) gradient (20%, 25%, 30%, 35%, 40%, 50%, and 60% in PBS) and centrifuged at 800 g for 30 min according to published protocol (Yoshikawa et al., 2009). Individual cell fraction was removed from the gradient, transferred to a conical test tube, and washed two times with PBS. 3μL of CellTracker™ CM-DiI (C7000, Invitrogen) labeling solution was added into 1 mL purified cell suspension to label the spermatogonia with red fluorescence. The cells were resuspended in 30 μL of L-15, and the number of red fluorescence positive spermatogonia was counted under a fluorescence microscopy using a hemocytometer. All of the cell counts are reported as mean ± SEM.

Spermatogonia obtained from the interfaces of the 30%–35% and 35%–40% Percoll fractions were combined, resuspended in 30 μL of L-15 medium, and immediately transplanted into zebrafish recipient larvae. Cell transplantations were performed using a glass micropipette needle (30 to 50 μm diameter borosilicate glass capillaries) under a stereomicroscope. Similar to the published methods using transplantation of trout PGCs into the peritoneal cavity (Takeuchi et al., 2003), the donor spermatogonia were transplanted into the abdominal cavity under the swim bladder close to the primitive gonads of newly hatched sterile zebrafish larvae. Three days after transplantation, all the recipients were examined by a fluorescence microscopy and the potential germline chimeras were identified based on the presence of red fluorescence positive cells in the gonad region. Three weeks after transplantation, 20 recipients (from 3 transplantations) were euthanized, and the colonization rates of the transplanted spermatogonia as well as its ability to proliferate in recipient gonads were evaluated. To determine if the transplanted spermatogonia were able to generate functional gametes in the recipients, the chimeric fish were raised to sexual maturity and paired with WT zebrafish. Three batches of eggs from each paired fish were collected, and the efficiency of fertilization was calculated.

### 4.6 Whole-mount in situ hybridization

Embryos were fixed with 4% paraformaldehyde (PFA)/PBS overnight at 4 °C. Digoxigenin (DIG)-labeled RNA probes were synthesized using T7 or T3 RNA polymerases (NEB, USA) with the DIG RNA Labeling Mix (Roche) according to the manufacturers’ protocols. WISH was performed essentially as described previously (Thisse et al., 2004).

### 4.7 Evaluation of mutation efficiency of fertile transplanted adults and genotyping of the mutated gametes

The spermatogonia transplanted zebrafish larvae were raised with great care and they usually became sexually matured at 2.5 months post transplantation (mpt). Adults of transplanted recipients were first crossed with WT zebrafish one by one to screen out those could produce sperm normally. Total DNA was isolated from the putative mutant embryos, and PCR was performed with certain primers (Tab. S1) which could amplify the target regions. The PCR products were subjected to sanger sequencing for sequence analysis to confirm that the sperm produced by recipients were target genes mutated or not.

To evaluate the mutation efficiencies of the gametes produced by the SSCs transplanted zebrafish, 8 to 11 hybrid embryos at 3 dpf were randomly selected for PCR amplification of the gRNA target sites and the amplicons were directly subject to sanger sequencing to analyze the mutation types and mutation efficiencies. Each fish was analyzed for three independent times. One positive transplanted zebrafish which could produce donor derived mutated gametes was mated with a target gene mutated *Gr* heterozygote. The incrossed F1 embryos were genotyped and phenotypically analyzed with a MV×10 microscope and used for further analysis.

### 4.8 Reverse-transcription PCR

Total RNA was isolated from testes of different samples by using Trizol method. The RNA was reverse-transcribed with PrimeScript™ RT reagent Kit (Takara) and RT-PCR was analyzed with gene specific primers listed in Table S1. *β-actin* was used as the internal control.

### 4.9 Histological analysis

For histological analysis, we dissected the intact testes from different samples at various developmental stages (40 dpf,2 mpf, 3 mpf,4 mpf) and fixed them in 4% PFA/PBS over night at 4 °C. After dehydration, samples were embedded in paraffin, and cut into slices with 4 μm thickness. Then the slices were stained with Hematoxylin and Eosin (H&E) according to standard protocols. The staging systems on spermatogenesis were identified as described previously (Schulz et al., 2010).

### 4.10 Sperm preparation for SEM and TEM

Milt from different samples were collected into a test tube and fixed with 100 μL 2.5% glutaraldehyde at 4 °C overnight. After centrifugation at 600 g/min for 2 min, the sperm pellet was washed with PBS for three times and resuspended in 100 μL PBS. The sperm suspension was then dropped onto a cell slide and left to dry naturally. Gradient alcohol series (10%, 30%, 50%, 70%, 80%, 90%, 95%, 100%, 100%) was used for dehydration and the slides were left to dry naturally. The samples were sputter-coated with gold (Hitachi, E-1010) and observed under SEM (Hitachi, S-4800). For TEM observation, samples were embedded in Epon812 resin and cutted into 70-80 nm slides with an ultrathin slicer after dehydration. The samples were then stained with uranium acetate and lead citrate and observed under TEM (Hitachi, HT7700).

### 4.11 Sperm motility test

Milt from different samples were collected into a test tube with Hank’s Buffer and chilled on ice. 0.5 μL milt was added into 500 μL H_2_O for activation and various kinetic parameters including VAP (average path velocity), VSL (straight line velocity), VCL (curvilinear velocity) and BCF (beating cross frequency) were measured using sperm class analyzer (SCA v5) for computer assisted semen analysis (CASA). Linearity (LIN) of sperm was the ratio of VSL to VCL.

### 4.12 EdU assay

Zebrafish larvae at 3 wpt were first incubated with 400 μM EdU for 20 hrs and then the testes from different samples were dissected and fixed in 4% PFA/PBS over night at 4 °C. After removing the 4% PFA/PBS, testes were neutralized with 2 mg/mL glycine solution at RT for 5min and then washed with 3% BSA/PBS twice. Next the testes were promoted for infiltration with 1% Triton X-100/PBS for about two hours at RT. Finally, the signaling of cell proliferation was detected with a Yefluor 594 EdU Imaging Kit (40276ES76, YEASEN, China) according to the manufacture’s instruction.

### 4.13 In situ hybridization on section

The complementary DNA (cDNA) was used for preparing probes, and the primers for PCR were listed in Table S1. Digoxigenin-labeled antisense RNA probes were synthesized by vitro transcription. For sections, the intact and fresh tissues were rinsed in ice-cold PBS for 10 minutes. Then the samples were embedded in a mixture of OCT and water at a ratio of 1:1 and sectioned at 10 μm thickness with freezing microtome (Leica). The mRNA detection by in situ hybridization was performed as previously described (Lauter et al., 2011; Rodriguez-Mari et al., 2005).

### 4.14 Immunofluorescence assay

Sections were prepared as in situ hybridization. After completely removing PFA with PBS, nonspecific antibodies were blocked with ice-cold PBST with BSA and DMSO (0.1% Triton-100, 1% BSA and 1% DMSO in PBS, pH7.4) for one hour at RT. Then, slides were incubated overnight at 4°C with primary antibodies diluted in blocking solution: anti-Vasa antibody, phosphohistone 3 (PH3) antibody (Cell Signaling Technology, 3377S), anti-Sycp3 antibody (ab150292). After washing, slides were incubated with secondary antibody (anti-Rabbit IgG, Alexa Fluor 488) diluted at 1:1000 in blocking solution overnight at 4 °C to detect primary antibodies. Following extensive washing, tissues were counterstained with DAPI and phalloidin (Molecular Probes) to label nuclei and cytoskeleton respectively and then the sections were photographed by a confocal microscopy.

### 4.15 Statistical analysis

Results were expressed as the mean ± SEM. Unpaired Student’s t-test was used to analyze the difference between two groups. One-way ANOVA statistics was used to analyze the difference among three independent samples.

## Acknowledgments

We sincerely thank Mrs. Ming Li at Institute of Hydrobiology, CAS for providing *gobiocypris rarus* embryos and we thank Kuoyu Li at the China Zebrafish Resource Center (CZRC) for zebrafish and *gobiocypris rarus* rearing. We also thank Mrs Fang Zhou and Yuan Xiao at analytical and testing center of Institute of Hydrobiology, CAS for providing technical support in confocal microscope, SEM and TEM analysis. This work was supported by National Natural Science Foundation of China (32025037 and 31721005), National Key R&D Project of China (2018YFA0801000 and 2018YFD0901205), Chinese Academy of Sciences (XDA24010108), and State Key Laboratory of Freshwater Ecology and Biotechnology (grant No. 2019FBZ05).

## Abbreviations

GSCs: germline stem cells
PGCs: primordial germ cells
SSCs: spermatogonial stem cells
OSCs: ovogenic stem cells
GSCT: germline stem cell transplantation
PGCT: primordial germ cell transplantation
SSCT: spermatogonial stem cell transplantation
OGCT: ovarian germ cell transplantation
CRISPR: clustered regularly interspaced short palindromic repeats
Cas9: CRISPR-associated 9
*Gr*: *gobiocypris rarus*
*Dr*: *danio rerio*
KO: knocked out
MO: morpholino
*dnd*: *dead end*
*buc*: *bucky ball*
GSI: gonadosomatic index
WT: wild type
dpf: days post fertilization
dpt: days post transplantation
wpt: weeks post transplantation
mpf: months post fertilization
SEM: scanning electron microscopy
TEM: transmission electron microscopy
WISH: whole-mount in situ hybridization
VAP: average path velocity
VSL: straight line velocity
VCL: curvilinear velocit
BCF: beat cross frequency
LIN: linearity
MHB: midbrain-hindbrain boundary.

## Notes

### Competing Interest Statement

The authors have declared no competing interest.

## References

Beer, R.L., Draper, B.W. (2013). Nanos3 maintains germline stem cells and expression of the conserved germline stem cell gene nanos2 in the zebrafish ovary. Developmental biology 374: 308–318

Bellaiche, J., Lareyre, J.J., Cauty, C., Yano, A., Allemand, I., Le Gac, F. (2014). Spermatogonial stem cell quest: Nanos2, marker of a subpopulation of undifferentiated a spermatogonia in trout testis. Biology of reproduction 90: 79

Brinster, R.L. (2002). Germline stem cell transplantation and transgenesis. Science 296: 2174–2176

Burgess, S., Reim, G., Chen, W.B., Hopkins, N., Brand, M. (2002). The zebrafish spiel-ohne-grenzen (spg) gene encodes the pou domain protein pou2 related to mammalian oct4 and is essential for formation of the midbrain and hindbrain, and for pre-gastrula morphogenesis. Development 129: 905–916

Chang, N.N., Sun, C.H., Gao, L., Zhu, D., Xu, X.F., Zhu, X.J., Xiong, J.W., Xi, J.J. (2013). Genome editing with rna-guided cas9 nuclease in zebrafish embryos. Cell research 23: 465–472

Cinalli, R.M., Rangan, P., Lehmann, R. (2008). Germ cells are forever. Cell 132: 559–562

Ciruna, B., Weidinger, G., Knaut, H., Thisse, B., Thisse, C., Raz, E., Schier, A.F. (2002). Production of maternal-zygotic mutant zebrafish by germ-line replacement. Proceedings of the National Academy of Sciences of the United States of America 99: 14919–14924

Eaton, R.C., Farley, R.D. (1974). Spawning cycle and egg-production of zebrafish, brachydanio-rerio, in laboratory. Copeia 195–204

Goto, R., Saito, T. (2019). A state-of-the-art review of surrogate propagation in fish. Theriogenology 133: 216–227

Gratacap, R.L., Wargelius, A., Edvardsen, R.B., Houston, R.D. (2019). Potential of genome editing to improve aquaculture breeding and production. Trends Genet 35: 672–684

Hammerschmidt, M., Pelegri, F., Mullins, M.C., Kane, D.A., Vaneeden, F.J.M., Granato, M., Brand, M., Furutaniseiki, M., Haffter, P., Heisenberg, C.P., et al. (1996). Dino and mercedes, two genes regulating dorsal development in the zebrafish embryo. Development 123: 95–102

Houston, R.D., Bean, T.P., Macqueen, D.J., Gundappa, M.K., Jin, Y.H., Jenkins, T.L., Selly, S.L.C., Martin, S.a.M., Stevens, J.R., Santos, E.M., et al. (2020). Harnessing genomics to fast-track genetic improvement in aquaculture. Nature Reviews Genetics

Ivell, R., Bathgate, R.a.D. (2002). Reproductive biology of the relaxin-like factor (rlf/insl3). Biology of reproduction 67: 699–705

Iwasaki-Takahashi, Y., Shikina, S., Watanabe, M., Banba, A., Yagisawa, M., Takahashi, K., Fujihara, R., Okabe, T., Valdez Jr, D.M., Yamauchi, A., et al. (2020). Production of functional eggs and sperm from in vitro-expanded type a spermatogonia in rainbow trout. Communications Biology 3:

Kimmel, C.B., Ballard, W.W., Kimmel, S.R., Ullmann, B., Schilling, T.F. (1995). Stages of embryonic-development of the zebrafish. Dev Dynam 203: 253–310

Kubota, H., Brinster, R.L. (2018). Spermatogonial stem cells†. Biology of Reproduction 99: 52–74

Lauter, G., Söll, I., Hauptmann, G. (2011). Multicolor fluorescent in situ hybridization to define abutting and overlapping gene expression in the embryonic zebrafish brain. Neural development 6: 10

Leal, M.C., Cardoso, E.R., Nobrega, R.H., Batlouni, S.R., Bogerd, J., Franca, L.R., Schulz, R.W. (2009). Histological and stereological evaluation of zebrafish (danio rerio) spermatogenesis with an emphasis on spermatogonial generations. Biology of reproduction 81: 177–187

Lehmann, R. (2012). Germline stem cells: Origin and destiny. Cell Stem Cell 10: 729–739

Li, M., Hong, N., Xu, H., Song, J., Hong, Y. (2016). Germline replacement by blastula cell transplantation in the fish medaka. Scientific reports 6:

Li, Q., Fujii, W., Naito, K., Yoshizaki, G. (2017). Application of dead end-knockout zebrafish as recipients of germ cell transplantation. Molecular reproduction and development 84: 1100–1111

Liang, X., Zha, J. (2016). Toxicogenomic applications of chinese rare minnow (gobiocypris rarus) in aquatic toxicology. Comparative biochemistry and physiology. Part D, Genomics & proteomics 19: 174–180

Lin, Q., Mei, J., Li, Z., Zhang, X., Zhou, L., Gui, J.F. (2017). Distinct and cooperative roles of amh and dmrt1 in self-renewal and differentiation of male germ cells in zebrafish. Genetics 207: 1007–1022

Lubzens, E., Young, G., Bobe, J., Cerda, J. (2010). Oogenesis in teleosts: How fish eggs are formed. General and comparative endocrinology 165: 367–389

Luo, S., Jin, S.Y., Su, L.X., Wang, J.W. (2017). Effect of water temperature on reproductive performance and offspring quality of rare minnow, gobiocypris rarus. Journal of thermal biology 67: 59–66

Moreno-Mateos, M.A., Vejnar, C.E., Beaudoin, J.D., Fernandez, J.P., Mis, E.K., Khokha, M.K., Giraldez, A.J. (2015). Crisprscan: Designing highly efficient sgrnas for crispr-cas9 targeting in vivo. Nat Methods 12: 982–988

Morita, T., Kumakura, N., Morishima, K., Mitsuboshi, T., Ishida, M., Hara, T., Kudo, S., Miwa, M., Ihara, S., Higuchi, K., et al. (2012). Production of donor-derived offspring by allogeneic transplantation of spermatogonia in the yellowtail (seriola quinqueradiata). Biology of reproduction 86:

Nasevicius, A., Ekker, S.C. (2000). Effective targeted gene ‘knockdown’ in zebrafish. Nature genetics 26: 216–220

Octavera, A., Yoshizaki, G. (2019). Production of donor-derived offspring by allogeneic transplantation of spermatogonia in chinese rosy bitterling. Biology of reproduction 100: 1108–1117

Ohta, H., Wakayama, T., Nishimune, Y. (2004). Commitment of fetal male germ cells to spermatogonial stem cells during mouse embryonic development. Biology of reproduction 70: 1286–1291

Okutsu, T., Suzuki, K., Takeuchi, Y., Takeuchi, T., Yoshizaki, G. (2006). Testicular germ cells can colonize sexually undifferentiated embryonic gonad and produce functional eggs in fish. Proceedings of the National Academy of Sciences of the United States of America 103: 2725–2729

Okutsu, T., Shikina, S., Kanno, M., Takeuchi, Y., Yoshizaki, G. (2007). Production of trout offspring from triploid salmon parents. Science 317: 1517

Perez-Cadahia, B., Drobic, B., Davie, J.R. (2009). H3 phosphorylation: Dual role in mitosis and interphase. Biochem Cell Biol 87: 695–709

Rodriguez-Mari, A., Yan, Y.L., Bremiller, R.A., Wilson, C., Canestro, C., Postlethwait, J.H. (2005). Characterization and expression pattern of zebrafish anti-mullerian hormone (amh) relative to sox9a, sox9b, and cyp19a1a, during gonad development. Gene expression patterns : GEP 5: 655–667

Saito, T., Fujimoto, T., Maegawa, S., Inoue, K., Tanaka, M., Arai, K., Yamaha, E. (2006). Visualization of primordial germ cells in vivo using gfp-nos1 3 ‘ utr mrna. Int J Dev Biol 50: 691–700

Sawatari, E., Shikina, S., Takeuchi, T., Yoshizaki, G. (2007). A novel transforming growth factor-beta superfamily member expressed in gonadal somatic cells enhances primordial germ cell and spermatogonial proliferation in rainbow trout (oncorhynchus mykiss). Developmental biology 301: 266–275

Schultemerker, S., Lee, K.J., Mcmahon, A.P., Hammerschmidt, M. (1997). The zebrafish organizer requires chordino. Nature 387: 862–863

Schulz, R.W., De França, L.R., Lareyre, J.J., Le Gac, F., Chiarini-Garcia, H., Nobrega, R.H., Miura, T. (2010). Spermatogenesis in fish. General and comparative endocrinology 165: 390–411

Slanchev, K., Stebler, J., De La Cueva-Mendez, G., Raz, E. (2005). Development without germ cells: The role of the germ line in zebrafish sex differentiation. P Natl Acad Sci USA 102: 4074–4079

Sun, Y., Zhu, Z. (2019). Designing future farmed fishes using genome editing. Science China. Life sciences 62: 420–422

Sun, Y.H., Zhang, B., Luo, L.F., Shi, D.L., Wang, H., Cui, Z.B., Huang, H.H., Cao, Y., Shu, X.D., Zhang, W.Q., et al. (2020). Systematic genome editing of the genes on zebrafish chromosome 1 by crispr/cas9. Genome research 30: 118–126

Tajima, A., Naito, M., Yasuda, Y., Kuwana, T. (1993). Production of germ-line chimera by transfer of primordial germ-cells in the domestic chicken (gallus-domesticus). Theriogenology 40: 509–519

Takeuchi, Y., Yoshizaki, G., Takeuchi, T. (2003). Generation of live fry from intraperitoneally transplanted primordial germ cells in rainbow trout. Biol Reprod 69: 1142–1149

Takeuchi, Y., Yoshizaki, G., Takeuchi, T. (2004). Biotechnology: Surrogate broodstock produces salmonids. Nature 430: 629–630

Thisse, B., Heyer, V., Lux, A., Alunni, V., Degrave, A., Seiliez, I., Kirchner, J., Parkhill, J.P., Thisse, C. (2004). Spatial and temporal expression of the zebrafish genome by large-scale in situ hybridization screening. Zebrafish:2nd Edition Genetics Genomics and Informatics 77: 505–519

Wang, J. (1992). Reproductive biology of gobiocypris rarus. Acta Hydrobiologica Sinica 16: 165–174

Wang, Y., Ge, W. (2004). Developmental profiles of activin βa, βb, and follistatin expression in the zebrafish ovary: Evidence for their differential roles during sexual maturation and ovulatory cycle1. Biol Reprod 71: 2056–2064

Weidinger, G., Stebler, J., Slanchev, K., Dumstrei, K., Wise, C., Lovell-Badge, R., Thisse, C., Thisse, B., Raz, E. (2003). Dead end, a novel vertebrate germ plasm component, is required for zebrafish primordial germ cell migration and survival. Current biology : CB 13: 1429–1434

Wong, T.T., Saito, T., Crodian, J., Collodi, P. (2011). Zebrafish germline chimeras produced by transplantation of ovarian germ cells into sterile host larvae. Biology of reproduction 84: 1190–1197

Yoshikawa, H., Ino, Y., Shigenaga, K., Katayama, T., Kuroyanagi, M., Yoshiura, Y. (2018). Production of tiger puffer takifugu rubripes from cryopreserved testicular germ cells using surrogate broodstock technology. Aquaculture 493: 302–313

Yoshikawa, H., Morishima, K., Fujimoto, T., Saito, T., Kobayashi, T., Yamaha, E., Arai, K. (2009). Chromosome doubling in early spermatogonia produces diploid spermatozoa in a natural clonal fish. Biology of reproduction 80: 973–979

Yoshikawa, H., Ino, Y., Kishimoto, K., Koyakumaru, H., Saito, T., Kinoshita, M., Yoshiura, Y. (2020). Induction of germ cell-deficiency in grass puffer by dead end 1 gene knockdown for use as a recipient in surrogate production of tiger puffer. Aquaculture 735385

Yoshikawa, H., Takeuchi, Y., Ino, Y., Wang, J., Iwata, G., Kabeya, N., Yazawa, R., Yoshizaki, G. (2017). Efficient production of donor-derived gametes from triploid recipients following intra-peritoneal germ cell transplantation into a marine teleost, nibe croaker (nibea mitsukurii). Aquaculture 478: 35–47

Yuan, L., Liu, J.G., Zhao, J., Brundell, E., Daneholt, B., Hoog, C. (2000). The murine scp3 gene is required for synaptonemal complex assembly, chromosome synapsis, and male fertility. Mol Cell 5: 73–83

Zhang, F.-H., Wang, H.-P., Huang, S.-Y., Xiong, F., Zhu, Z.-Y., Sun, Y.-H. (2016). A comparison of the knockout efficiencies of two codon-optimized cas9 coding sequences in zebrafish embryos. Yi chuan = Hereditas 38: 144–154

Zhang, F., Li, X., He, M., Ye, D., Xiong, F., Amin, G., Zhu, Z., Sun, Y. (2020). Efficient generation of zebrafish maternal-zygotic mutants through transplantation of ectopically induced and cas9/grna targeted primordial germ cells. Journal of genetics and genomics = Yi chuan xue bao 47: 37–47

