## Supplementary material for "Surrogate production of genome edited sperm from a different subfamily by spermatogonial stem cell transplantation": Figure S

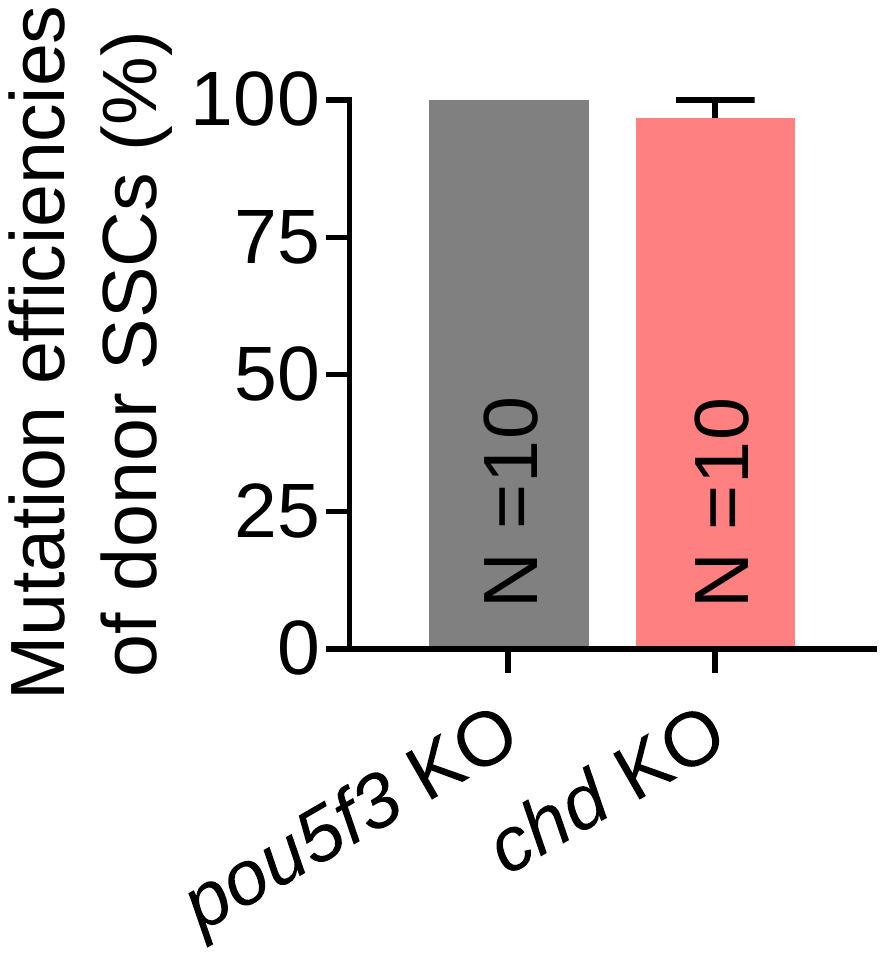

**Figure S1 High mutation efficiencies of SSCs isolated from testes of *pou5f3* and *chd* knocked out F0 adult males.**

The mutation efficiencies reach as high as 100.0% and 96.7% for *pou5f3* and *chd* KO SSCs respectively. N, number of clones sequenced. The experiment was replicated for three times.

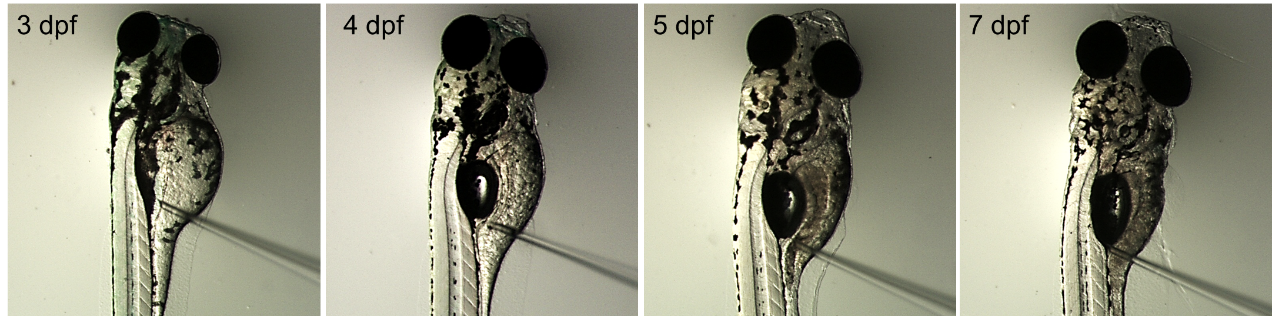

**Figure S2 Different stages of zebrafish larvae were utilized as host for SSCT.**

To optimize the zebrafish host for SSCT, 3, 4, 5 and 7 dpf zebrafish larvae were utilized as recipients. The donors SSCs were transplanted into the abdominal cavity between the swim bladder and the gut.

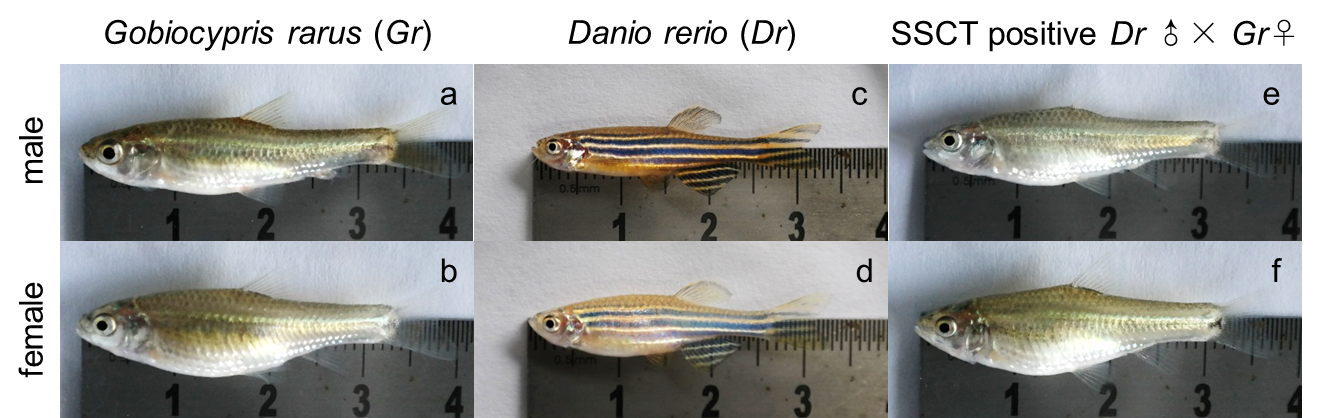

**Figure S3 The progeny of SSCT positive zebrafish male and rare minnow female could grow to adulthood normally.**

The phenotype of WT rare minnow male and female was shown in a and b; the phenotype of WT zebrafish male and female was shown in c and d; the progeny of SSCT positive zebrafish male and rare minnow female could grow to adulthood normally, and both male (e) and female (f) were obtained.

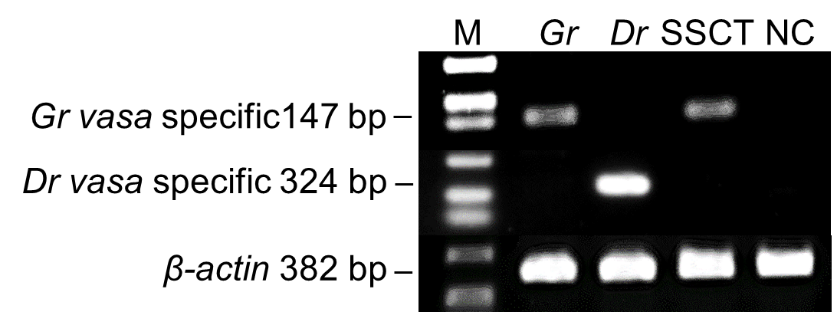

**Figure S4 *Gr* *vasa* but not *Dr* *vasa* was expressed in SSCT positive *Dr* testis**

RT-PCR using *Gr* and *Dr* specific *vasa* primers showed that they specifically expressed in *Gr* and *Dr* testes respectively, while *Gr* *vasa* but not *Dr* *vasa* expressed in SSCT positive *Dr* testis. Neither *Gr* *vasa* nor *Dr vasa* was detected in *dnd* MO injected *Dr* testis. M, DS2000 DNA maker; *Gr*, amplicons from *gobiocypris rarus* testes cDNA; *Dr*, amplicons from zebrafish testes cDNA; SSCT, amplicons from SSCT positive zebrafish testes cDNA; NC, amplicon from *dnd* MO injected zebrafish testes cDNA. *β-actin* was utilized as an internal reference.

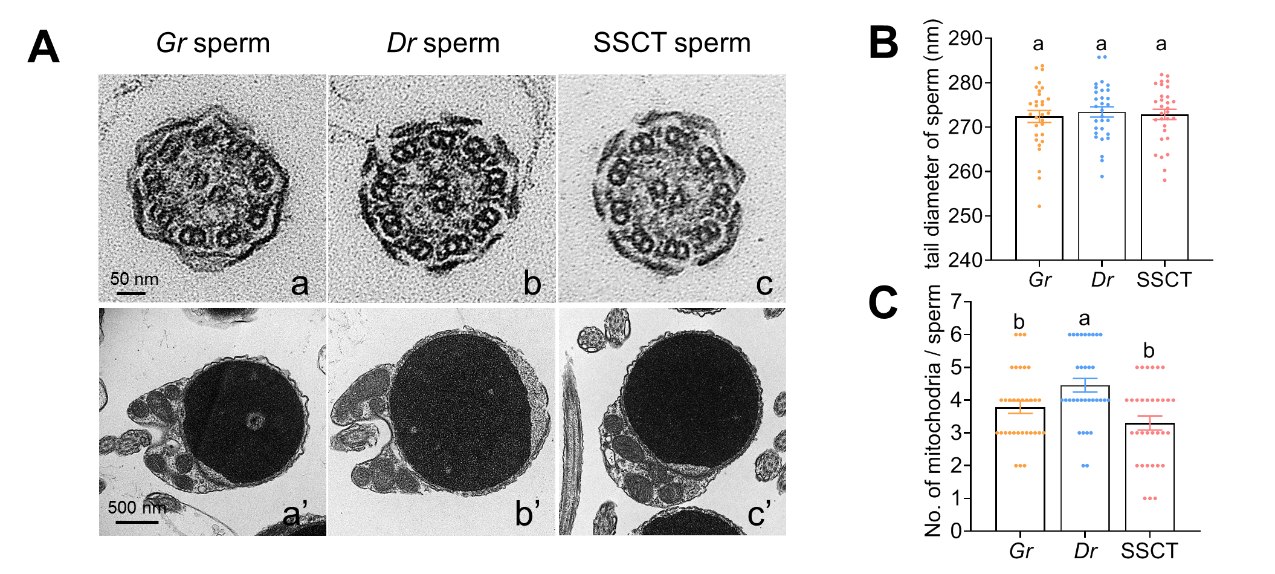

**Figure S5 Inner structure observation of sperms from *Gr*, *Dr* and SSCT positive *Dr* testis by TEM**

**A**: Sections of tails and heads of sperms from *Gr*, *Dr* and SSCT positive *Dr* testis. Note that the center of the flagella of three kinds of sperm is the axial filament with a typical “9×2+2” microtubule structure (a,b,c). The membrane of head showed serrated phenotype both in *Gr* (a’) and SSCT sperm (c’) while it’s smooth in *Dr* sperm (b’). *Dr* sperm possesses overt cytoplasm at the front of the head, but it’s not seen neither in *Gr* nor SSCT sperm head.

**B**: There was no significant differences in tail diameter among *Gr*, *Dr* and SSCT sperm.

**C**: Statistics of mitochondria number in *Gr*, *Dr* and SSCT sperm. The mitochondria number was slightly reduced in SSCT sperm compared to *Dr* sperm.

Supplemental Table 1. Primers used in the present study

| Primers | Sequences (5’ to 3’) | usage |
| --- | --- | --- |
| *chd*-Cas9-F4 | GCGGTTCCGCTCTGACTC | for amplifying target sites |
| *chd*-Cas9-R4 | GCCGTGGCGATGTTGATG |  |
| *pou5f3-*Cas9-F2 | CGAACGATTTGCAAGGT |  |
| *pou5f3-*Cas9-R2 | TGCTCCTGGCTGTGCTTTG |  |
| *chd-*gRNA-F4 | TGTAATACGACTCACTATAggagcgcgg  cggaggccgggGTTTTAGAGCTAGAAAT | for amplifying gRNA templates |
| *pou5f3*-gRNA-F2 | TGTAATACGACTCACTATAgggacgtca  tccgtgcccacGTTTTAGAGCTAGAAAT |  |
| gRNA-RP | AAAAAAAGCACCGACTCGGTGCCAC |  |
| *Gr*-*chd*-F1 | GCGGGCAGCTCCAGATCTC | for amplifying *chd* and *pou5f3* partial CDS and genome sequences |
| *Gr*-*chd*-R1 | GTGGCGATGTTGATGAACAG |  |
| *Gr*-*chd*-F2 | CACATGCTGCTGCAGAAC |  |
| *Gr*-*chd*-R2 | CATCTTGGTGCTGACCTG |  |
| *Gr*-*pou5f3*-F1 | GGTCAACAGGGCCATGTA |  |
| *Gr*-*pou5f3*-R1 | CTGACTGAACATTTTGCC |  |
| *Gr*-*pou5f3*-F2 | ACACTGGTACCCGTTTGC |  |
| *Gr*-*pou5f3*-R2 | TGTCTACGGTTGCAGAACC |  |
| *Gr*-*chd-*spe-F | GCGGTTCCGCTCTGACTC | for amplifying partial fragments of *chd* genome |
| *Gr*-*chd-*spe-R | GCCGTGGCGATGTTGATG |  |
| *Dr*-*chd-*spe-F | ATCCATCAATCCATTATCTT |  |
| *Dr*-*chd-*spe-R | TGGTCTGTGAACACTGCC |  |
| *Gr*-*nanos2*-pro-F | GGACCCTGTCGCAGCTGCTG | for amplifying probe templates |
| *Gr*-*nanos2*-pro-R | TAATACGACTCACTATAGGTGTTCCTCCGCGGACAGTAGT |  |
| *Gr*-*vasa*-pro-F | CAACATTTCTCTGTCAGGAG |  |
| *Gr*-*vasa*-pro-R | TAATACGACTCACTATAGGCAAAAGTTCTTCCACGAGG |  |
| *Dr*-*vasa*-pro-F | GTCTTGCAGCTCAGGATT |  |
| *Dr*-*vasa*-pro-R | TAATACGACTCACTATAGGAAATTAATGCCTGTTGCATA |  |
| *Dr*-*insl3-*pro-F | GTGGTCGTGAGTTCGTTC |  |
| *Dr*-*insl3-*pro-R | TAATACGACTCACTATAGGGGTCGGCTCCAGAGTAAAT |  |
| *Dr*-*gsdf-*pro-F | GACACACTCGACCCCGCAGC |  |
| *Dr*-*gsdf-*pro-R | TAATACGACTCACTATAGGGCTGCCAGAGCCAAACCCGCA |  |
| *Gr*-*insl3-*pro-F | ACATAAGCCCACCAGCAGAGC |  |
| *Gr*-*insl3-*pro-R | TAATACGACTCACTATAGGTTCATGTTAGCTCAGGTCT |  |
| *Gr*-*gsdf-*pro-F | TCTTCATGAAAGATCTGGGCT |  |
| *Gr*-*gsdf-*pro-R | TAATACGACTCACTATAGGTTTGGAGGGAACACTCAGT |  |
| *Gr*-*pou5f3-*pro-F | GGTCAAAACGGAGAAAGA |  |
| *Gr*-*pou5f3-*pro-R | TAATACGACTCACTATAGGCTCAAAGCGGCAGATAGT |  |
| *Gr*-*chd-*pro-F | GATGGGAGGAATCGCCATG |  |
| *Gr*-*chd-*pro-R | TAATACGACTCACTATAGGCACTCCTTACAGCAGTC |  |
| *Gr*-*gsc-*pro-F | GGTTGTGTTCTCCAACTTG |  |
| *Gr*-*gsc-*pro-R | TAATACGACTCACTATAGGGCTCGTCTGTTTTTGAACC |  |
| *Gr*-*eve1-*pro-F | CCGGACTGCTTTCACGAGG |  |
| *Gr*-*eve1-*pro-R | TAATACGACTCACTATAGGTCTGGTCTCCAGTGCAGGCA |  |
| *Gr*-*krox20-*pro-F | AGCTTCGCGCAACCAGCT |  |
| *Gr*-*krox20-*pro-R | TAATACGACTCACTATAGGGTTCCTGATAGTGTTCAGG |  |
| *Gr*-*wnt1-*pro-F | TCCTCTCTCACTGGCACAG |  |
| *Gr*-*wnt1-*pro-R | TAATACGACTCACTATAGGTGTTGTGCAGGTTGGTCAG |  |
| *Gr*-*vasa-*spe-F | GGGAAAGGGTTCTGGTTTTC | for RT-PCR analysis |
| *Gr*-*vasa-*spe-R | GCAGACGATTAAGTATTTCC |  |
| *Dr*-*vasa-*spe-F | GTGCAGAAGTGGGTGGGTTTGT |  |
| *Dr*-*vasa-*spe-R | CACCGCTCTGTTACAGTCAGTC |  |
| *β-actin*-RT-F | CGTTGACAACGGCTCCGGTATG |  |
| *β-actin*-RT-R | GATGGCCACATACATGGCAGG |  |

Supplemental Table 2. mutation types of *pou5f3* mutated *Gr* sperm derived from SSCT positive *Dr* males

| Allele | Target sequence (#1) | Test No. | | |
| --- | --- | --- | --- | --- |
|  |  | 8 | 8 | 8 |
| WT | CTCACTCCCGGGACGTCATCCGTGCCCACCGGGGTGAATTATTA | 0 | 0 | 0 |
| -1bp | CTCACTCCCGGGACGTCATCCGTGCC-ACCGGGGTGAATTATTA | 2 | 5 | 4 |
| -8+6bp | CTCACTCCCGGGACGTCATAATAAT--ACCGGGGTGAATTATTA | 6 | 3 | 4 |
| Allele | Target sequence (#2) | Test No. | | |
|  |  | 11 | 11 | 11 |
| WT | CTCACTCCCGGGACGTCATCCGTGCCCACCGGGGTGAATTATTA | 0 | 0 | 0 |
| -4bp | CTCACTCCCGGGACGTCATCCGTG**CC**----GGGGTGAATTATTA | 5 | 7 | 6 |
| -4+9bp | CTCACTCCCGGGACGTCATCCGTGGGTAATTAACCGGGGTGAAT  TATTA | 6 | 4 | 5 |
| Allele | Target sequence (#3) | Test No. | | |
|  |  | 10 | 10 | 8 |
| WT | CTCACTCCCGGGACGTCATCCGTGCCCACCGGGGTGAATTATTA | 0 | 0 | 0 |
| -4bp | CTCACTCCCGGGACGTCATCCGTG**CC**----GGGGTGAATTATTA | 3 | 3 | 1 |
| -10bp | CTCACTCCCGGGACGTCATC----------GGGGTGAATTATTA | 2 | 4 | 3 |
| -12bp | CTCACTCCCGGGACGTCATCC**GTG**------------AATTATTA | 4 | 2 | 3 |
| -11+5bp | CTCACTCCCGGGACGTAATTC------ACCGGGGTGAATTATTA | 1 | 1 | 0 |
| -5bp | CTCACTCCCGGGACGTCATCCGTGC-----GGGGTGAATTATTA | 0 | 0 | 1 |
| Allele | Target sequence (#4) | Test No. | | |
|  |  | 8 | 8 | 8 |
| WT | CTCACTCCCGGGACGTCATCCGTGCCCACCGGGGTGAATTATTA | 0 | 0 | 0 |
| -10bp | CTCACTCCCGGGACGT**CA**----------CCGGGGTGAATTATTA | 4 | 5 | 5 |
| -2+21bp | CTCACTCCCGGGACGTCATCCGTGCCGAGGTGTGAATTATTACA  CGCCCGGGGTGAATTATTA | 4 | 3 | 3 |
| Allele | Target sequence (#5) | Test No. | | |
|  |  | 8 | 8 | 8 |
| WT | CTCACTCCCGGGACGTCATCCGTGCCCACCGGGGTGAATTATTA | 0 | 0 | 0 |
| -8+6bp | CTCACTCCCGGGACGTCATAATAAT--ACCGGGGTGAATTATTA | 3 | 3 | 4 |
| -13bp | CTCACTCCCGGGACGTCATCCGT-------------AATTATTA | 4 | 1 | 2 |
| -6+7bp | CTCACTCCCGGGACGTCATCAATAATTCACCGGGGTGAATTATT  A | 1 | 4 | 2 |
| Allele | Target sequence (#6) | Test No. | | |
|  |  | 8 | 8 | 8 |
| WT | CTCACTCCCGGGACGTCATCCGTGCCCACCGGGGTGAATTATTA | 0 | 0 | 0 |
| -1bp | CTCACTCCCGGGACGTCATCCGTGCC-ACCGGGGTGAATTATTA | 3 | 2 | 2 |
| -8+6bp | CTCACTCCCGGGACGTCATAATAAT--ACCGGGGTGAATTATTA | 5 | 6 | 6 |
| Allele | Target sequence (#7) | Test No. | | |
|  |  | 8 | 8 | 8 |
| WT | CTCACTCCCGGGACGTCATCCGTGCCCACCGGGGTGAATTATTA | 0 | 0 | 0 |
| -4bp | CTCACTCCCGGGACGTCATCCGTG**CC**----GGGGTGAATTATTA | 4 | 5 | 3 |
| -9+7bp | CTCACTCCCGGGACGTCATCCGTGCC--TGTGGGAGAATTATTA | 0 | 1 | 1 |
| -9bp | CTCACTCCCGGGACGTCAT**CCG**---------GGGTGAATTATTA | 2 | 2 | 3 |
| -8bp | CTCACTCCCGGGACGTCATCCGTG--------GGTGAATTATTA | 1 | 0 | 0 |
| +17bp | CTCACTCCCGGGACGTCATCCGTGCCACCCCGGTGTAATAATTC  ACCGGGGTGAATTATTA | 1 | 0 | 1 |
| Allele | Target sequence (#8) | Test No. | | |
|  |  | 8 | 8 | 8 |
| WT | CTCACTCCCGGGACGTCATCCGTGCCCACCGGGGTGAATTATTA | 0 | 0 | 0 |
| -7bp&1bp trans | CTCACTCCCGGGACGACATCCGTG-------GGGTGAATTATTA | 3 | 5 | 3 |
| -3+9bp | CTCACTCCCGGGACGTCATCCGTGCCGGATGACGTCGGGGTGAA  TTATTA | 5 | 3 | 5 |
| Allele | Target sequence (#9) | Test No. | | |
|  |  | 10 | 10 | 10 |
| WT | CTCACTCCCGGGACGTCATCCGTGCCCACCGGGGTGAATTATTA | 0 | 0 | 0 |
| -4bp | CTCACTCCCGGGACGTCATCCGT----ACCGGGGTGAATTATTA | 0 | 1 | 0 |
| -3+1bp | CTCACTCCCGGGACGTCATCCGTGT--ACCGGGGTGAATTATTA | 4 | 4 | 3 |
| -7bp&1bp trans | CTCACTCCCGGGACGACATCCGTG-------GGGTGAATTATTA | 2 | 2 | 1 |
| -3+11bp | CTCACTCCCGGGACGTCATCCGTCATCCGTAATTCACCGGGGTG  AATTATTA | 1 | 2 | 3 |
| -3+22bp | CTCACTCCCGGGACGTCATCCGTGGTGAATTATTACACGCCCTG  GAACCGGGGTGAATTATT | 3 | 1 | 3 |
| Allele | Target sequence (#10) | Test No. | | |
|  |  | 8 | 8 | 8 |
| WT | CTCACTCCCGGGACGTCATCCGTGCCCACCGGGGTGAATTATTA | 0 | 0 | 0 |
| -7bp&1bp trans | CTCACTCCCGGGACGACATCCGTG-------GGGTGAATTATTA | 7 | 5 | 6 |
| -4bp | CTCACTCCCGGGACGTCATCCGTG**CC**----GGGGTGAATTATTA | 1 | 2 | 1 |
| -9bp | CTCACTCCCGGGACGTCAT**CCG**---------GGGTGAATTATTA | 0 | 1 | 1 |

WT, wild type; bp, base pair; the sequences underlined represent target site, sequences in blue represent PAM, the red dotted line and sequences represent loss or insertion of bases, sequences in green represent base transition or transversion, sequences in bold show DNA repair by microhomology-mediated end joining (MMEJ).

Supplemental Table 3. mutation types of *chd* mutated *Gr* sperm derived from SSCT positive *Dr* males

| Allele | Target sequence (#1) | Test No. | | |
| --- | --- | --- | --- | --- |
|  |  | 9 | 9 | 9 |
| WT | AGCGGAGGGGGAGCGCGGCGGAGGCCGGGCGGGGGGCAGCTGCA | 0 | 0 | 0 |
| +2bp | AGCGGAGGGGGAGCGCGGCGGAGGCCGGGGGCGGGGGGCAGCTG  CA | 5 | 3 | 4 |
| -12bp | AGCGGAGGGGGAGCGCGGCGGA**GG**------------CAGCTGCA | 4 | 6 | 5 |
| Allele | Target sequence (#2) | Test No. | | |
|  |  | 10 | 10 | 8 |
| WT | AGCGGAGGGGGAGCGCGGCGGAGGCCGGGCGGGGGGCAGCTGCA | 0 | 0 | 0 |
| -5bp | AGCGGAGGGGGAGCGCGGCGGA**GG**-----CGGGGGGCAGCTGCA | 5 | 4 | 3 |
| -8+3bp | AGCGGAGGGGGAGCGCGGCGGAGGCCCCC-----GGCAGCTGCA | 2 | 1 | 2 |
| -14bp | AGCGGAGGGGGAGC**GCG**--------------GGGGGCAGCTGCA | 3 | 5 | 3 |
| Allele | Target sequence (#3) | Test No. | | |
|  |  | 8 | 8 | 8 |
| WT | AGCGGAGGGGGAGCGCGGCGGAGGCCGGGCGGGGGGCAGCTGCA | 0 | 0 | 0 |
| -12bp | AGCGGAGGGGGAGCGCGGCGGA**GG**------------CAGCTGCA | 6 | 7 | 5 |
| -34bp | AGCGGAGGGGGAGCGCGG--------------------------  -------- | 2 | 1 | 3 |
| Allele | Target sequence (#4) | Test No. | | |
|  |  | 9 | 9 | 9 |
| WT | AGCGGAGGGGGAGCGCGGCGGAGGCCGGGCGGGGGGCAGCTGCA | 0 | 0 | 0 |
| +2bp | AGCGGAGGGGGAGCGCGGCGGAGGCCGGGGGCGGGGGGCAGCTG  CA | 6 | 5 | 5 |
| -27bp | A**GCGG**---------------------------GGGGCAGCTGCA | 3 | 4 | 4 |
| Allele | Target sequence (#5) | Test No. | | |
|  |  | 9 | 9 | 9 |
| WT | AGCGGAGGGGGAGCGCGGCGGAGGCCGGGCGGGGGGCAGCTGCA | 0 | 0 | 0 |
| -11bp | AGCGGAGGGGGAGCGC**GGCGG**-----------GGGGCAGCTGCA | 4 | 5 | 6 |
| -14bp | AGCGGAGGGGGAGC**GCG**--------------GGGGGCAGCTGCA | 4 | 4 | 3 |
| 4bp trans | AGCGGAGGGGGAGCGCGGCGGAGGCAGGTGAGGGGGCAGCTGCA | 1 | 0 | 0 |
| Allele | Target sequence (#6) | Test No. | | |
|  |  | 9 | 9 | 9 |
| WT | AGCGGAGGGGGAGCGCGGCGGAGGCCGGGCGGGGGGCAGCTGCA | 0 | 0 | 0 |
| -27bp | A**GCGG**---------------------------GGGGCAGCTGCA | 5 | 6 | 6 |
| -1+4bp | AGCGGAGGGGGAGCGCGGCGGAGGCAGCTGGGCGGGGGGCAGCT  GCA | 4 | 3 | 3 |
| Allele | Target sequence (#7) | Test No. | | |
|  |  | 10 | 10 | 10 |
| WT | AGCGGAGGGGGAGCGCGGCGGAGGCCGGGCGGGGGGCAGCTGCA | 0 | 0 | 0 |
| -11bp | AGCGGAGGGGGAGCGC**GGCGG**-----------GGGGCAGCTGCA | 4 | 5 | 7 |
| -20bp | AGCGGA**GGG**--------------------CGGGGGGCAGCTGCA | 5 | 4 | 2 |
| -1+4bp | AGCGGAGGGGGAGCGCGGCGGAGGCAGCTGGGCGGGGGGCAGCT  GCA | 1 | 0 | 1 |
| -1+13bp | AGCGGAGGGGGAGCGCGGCGGAGGCAGCTGCAGCCGCGGGGCGG  GGGGCAGCTGCA | 0 | 1 | 0 |
| Allele | Target sequence (#8) | Test No. | | |
|  |  | 9 | 9 | 8 |
| WT | AGCGGAGGGGGAGCGCGGCGGAGGCCGGGCGGGGGGCAGCTGCA | 0 | 0 | 0 |
| -10bp | AGCGGAGGGGGAGCGCGGCGGA**GG**----------GGCAGCTGCA | 6 | 7 | 5 |
| -26bp | AGCGGAGGGGGAGCGC--------------------------CA | 2 | 2 | 3 |
| -5+4bp | AGCGGAGGGGGAGCGCGGCGGAGGCCAGCT-GGGGGCAGCTGCA | 1 | 0 | 0 |
| Allele | Target sequence (#9) | Test No. | | |
|  |  | 8 | 8 | 8 |
| WT | AGCGGAGGGGGAGCGCGGCGGAGGCCGGGCGGGGGGCAGCTGCA | 0 | 0 | 0 |
| -12bp | AGCGGAGGGGGAGCGCGGCGGA**GG**------------CAGCTGCA | 3 | 2 | 4 |
| -27bp | AGCGGAGGGGGAGC**GC**---------------------------A | 1 | 3 | 1 |
| -4bp | AGCGGAGGGGGAGCGCGGCGGA**GG**----GCGGGGGGCAGCTGCA | 4 | 3 | 2 |
| -7+17bp | AGCGGAGGGGGAGCGCGGCGGAGGGGCAGCTGCAGCCGCGTGGG  GGCAGCTGCA | 0 | 0 | 1 |

WT, wild type; bp, base pair; the sequences underlined represent target site, sequences in blue represent PAM, the red dotted line and sequences represent loss or insertion of bases, sequences in green represent base transition or transversion, sequences in bold show DNA repair by microhomology-mediated end joining (MMEJ).
